# Life history and cancer in birds: clutch size predicts cancer

**DOI:** 10.1101/2023.02.11.528100

**Authors:** Stefania E. Kapsetaki, Zachary Compton, Jordyn Dolan, Valerie K. Harris, Shawn M. Rupp, Elizabeth G. Duke, Tara M. Harrison, Selin Aksoy, Mathieu Giraudeau, Orsolya Vincze, Kevin J. McGraw, Athena Aktipis, Marc Tollis, Amy M. Boddy, Carlo C. Maley

**Affiliations:** Arizona Cancer Evolution Center, Arizona State University, Tempe, AZ, USA; Department of Psychology, Arizona State University, Tempe, AZ, USA; Department of Anthropology, University of California Santa Barbara, CA, USA; Center for Biocomputing, Security and Society, Biodesign Institute, Arizona State University, Tempe, AZ, USA; School of Life Sciences, Arizona State University, Tempe, AZ, USA; Department of Clinical Sciences, North Carolina State University, Raleigh, NC, 27607, USA; Exotic Species Cancer Research Alliance, North Carolina State University, Raleigh, NC, 27607, USA; UMR IRD, CREEC, Université de Montpellier, 224-CNRS 5290 Montpellier, France; Centre de Recherche en Écologie Et Évolution de La Sante (CREES), Montpellier, France; Littoral Environnement Et Sociétés (LIENSs), UMR 7266, CNRS-La Rochelle Université, La Rochelle, France; Evolutionary Ecology Group, Hungarian Department of Biology and Ecology, Babeș-Bolyai University, Cluj-Napoca, Romania; Institute of Aquatic Ecology, Centre for Ecological Research, Debrecen, Hungary; School of Informatics, Computing, and Cyber Systems, Northern Arizona University, PO Box 5693, Flagstaff, AZ 8601, USA

## Abstract

Cancer is a disease that affects nearly all multicellular life, including birds. However, little is known about what factors explain the variance in cancer prevalence among species. Litter size is positively correlated with cancer prevalence in managed species of mammals, and larger body size, but not incubation or nestling period, is linked to tumor prevalence in wild birds. Also, birds that produce more elaborate sexual traits are expected to have fewer resources for cancer defenses and thus higher cancer prevalence. In this study, we examined whether cancer prevalence is associated with a wide variety of life history traits (clutch size, incubation length, body mass, lifespan, and the extent of sexual dimorphism) across 108 species of managed birds in 25 different zoological facilities, sanctuaries, and veterinary clinics. We found that clutch size was positively correlated with cancer and neoplasia (both benign and malignant) prevalence, even after controlling for body mass. Cancer prevalence was not associated with incubation length, body mass, lifespan, or sexual dimorphism. The positive correlations of clutch size with cancer prevalence and neoplasia prevalence suggest that there may be life-history trade-offs between reproductive investment and somatic maintenance (in the form of cancer prevention mechanisms) in managed birds.

## Introduction

Nearly all multicellular organisms are susceptible to neoplastic disease^1, 2^. Neoplasia is a disease consisting of uncontrolled cell division and growth, resulting ultimately in the formation of a tumor, as well as invasion or metastasis in case of malignant neoplasia (aka cancer)^3, 4^. Over the past few decades, cancer research has focused on identifying different molecular pathways, hallmarks, and control mechanisms of cancer – all with the ultimate aim of improving cancer treatment^5, 6^. Evolutionary biology has also been an important component of cancer research over the last 50 years^3, 7^. The ecological conditions under which organisms evolved have shaped their responses to various diseases, including cancer^8, 9^. Understanding why organisms differ in their ability to suppress cancer, as well as how they respond to neoplastic expansion, is a central question in comparative cancer research.

In general, life history trade-offs govern how organisms allocate time and resources to fitness components such as growth, self (or somatic)-maintenance, and reproduction^10, 11^. Somatic maintenance can include tumor suppression mechanisms such as cell cycle control and DNA damage repair. These trade-offs may help explain the variation in cancer prevalence across species. For example, long-lived species that invest in somatic maintenance over reproduction likely evolved enhanced mechanisms to suppress or evade cancer during their relatively long lifespans compared to short-lived species that invest heavily in reproductive effort rather than somatic maintenance^12^. Peto’s paradox predicts that bigger-longer lived animals would not be more vulnerable to cancer^13–15^. Utilizing this life history tradeoff approach can both give us insight into the basic biology and origins of cancer and also provide opportunities to discover either universal or novel mechanisms of cancer suppression that could have clinical applications to humans.

Birds (class Aves) represent a diverse vertebrate clade with considerable variation in life-history characteristics. This makes birds a suitable system for investigating the correlation between cancer risk and certain phenotypic traits, such as body mass and lifespan. Double-barred finches weigh on average just 9.5 grams, whereas greater rheas weigh on average 23 kilograms. Gouldian finches live on average up to six months, whereas salmon-crested cockatoos live on average up to 65 years (supplementary data). Birds also have a ZW genetic sex determination system, with females as the heterogametic sex, and therefore can also shed light on possible sex biases in health outcomes. For instance, female birds may be more susceptible to deleterious mutations promoting cancer development, whereas male birds may be protected by non-mutant versions of those alleles on their extra Z chromosome. This is known in humans as the two-X chromosome theory of cancer protection^16^. If this two-chromosome theory is correct, we would expect female birds to have higher cancer prevalence than male birds.

Cancer prevalence in birds has been an area of ongoing study. Previous work reports that birds have on average the lowest cancer prevalence amongst vertebrates^2, 17, 18^. Within birds, there is much variation in cancer prevalence which may be explained by some phenotypic traits. For instance, Møller et al. surveyed free-living Eurasian birds post-mortem and found that, when analyzing at least 20 individuals per species, larger body size was correlated with tumor prevalence^19^, while neither incubation nor nestling time were correlated with tumor prevalence^19^. Separate studies have reported neoplasms (benign and malignant tumors combined) in bird species, either free-living or in human care^2, 20–25^, but the prevalence of malignancy itself has not been measured before across bird species.

Clutch size could also be an important factor influencing the amount of energy devoted to somatic maintenance, including immune function, given the energetic trade-off between maintenance of a particular species’ own body versus its offspring^26, 27^. There may also be a trade-off between reproductive investments and somatic maintenance^28^ such that sexually dimorphic or dichromatic species experience increased cancer prevalence^29^ due to the somatic maintenance costs incurred by growing and maintaining these exaggerated morphological traits^30–33^. However, there has not been a study investigating the relationship between reproductive or sexually selected traits and cancer prevalence in birds.

To investigate the relationship between life history and cancer risk in birds, we combined trait-rich life-history databases with cancer prevalence data from veterinary records of 108 bird species under managed care. We hypothesized that the incredible diversity of life-history strategies observed across the class Aves can explain taxonomic differences in cancer risk in birds, due to the evolutionary trade-offs between growth, reproduction, and somatic maintenance. We test Peto’s paradox (under the expectation that body mass does not explain variation in cancer prevalence) in birds, and investigate whether malignancy prevalence or neoplasia prevalence is correlated with other avian traits such as incubation length, clutch size, and degree of sexual dimorphism and dichromatism. We also test for sex differences in cancer prevalence in birds, e.g., whether female birds (ZW sex chromosomes) have higher cancer prevalence than male birds (ZZ sex chromosomes). This study is the first to examine a wide range of life history traits in birds in order to predict cancer prevalence.

## Methods

### Cancer data from managed populations of birds

To collect avian cancer records, we collaborated with numerous zoological facilities, sanctuaries, and veterinary clinics. The data represent over 25 years of pathology records from 25 different institutions using 5,499 individual necropsies, including descriptions of age at death of 1287 individuals from 51 species, and malignancies and benign tumors across 108 bird species across 24 different avian orders managed under human care^34^. We measured malignancy prevalence and neoplasia prevalence (benign and malignant tumor) for each species by dividing the total number of necropsies reporting malignancies (or neoplasms) by the total number of necropsies available for that species (supplementary data); a measurement also used in previous studies^9, 35^.

### Life-history data

We assembled life-history variables from multiple published resources, including AnAge^36^ and the Amniote Life History Database^37^. The collected life-history variables included species averages of adult body mass (g), lifespan (months), incubation length (months), clutch size (number of offspring)^36, 37^, presence and degree of sexual plumage dichromatism (plumage brightness and plumage hue)^38^, and sexual size dimorphism (mass and tail size)^39^.

### Data filtering

We only included bird species for which we had at least 20 necropsies in our analysis. For analyses comparing female and male malignancy prevalence or neoplasia prevalence, as well as sex bias regressions, we used species with at least 10 necropsy records per sex. We present the neoplasia and malignancy prevalence of 108 bird species (supplementary data). We were not able to find data on every life-history variable for every species, so in the life-history analyses, the number of species is less than 108 (body mass correlations: 100 species; lifespan correlations: 59 species; body mass x lifespan correlations: 57 species; incubation length correlations: 34 species; clutch size correlations including domesticated/semi-domesticated species: 51 species; clutch size correlations excluding domesticated/semi-domesticated species: 45 species; dimorphism in brightness correlations: 18 species; dimorphism in hue correlations: 24 species; dimorphism in mass correlations: 47 species; dimorphism in tail size correlations: 34 species; sex differences in neoplasia prevalence: 31 species). We removed all necropsies from birds that had lived in the wild. We excluded chickens (*Gallus gallus*) from the analyses because as a largely domesticated agricultural species they have been selected for egg laying and frequently develop ovarian cancer^40^. We only included chickens (*Gallus gallus*) in Table 1 and in the Supp. Fig. 3 illustration of normalized frequency of the species’ age at death as a percentage of the species lifespan.

**Table 1.**
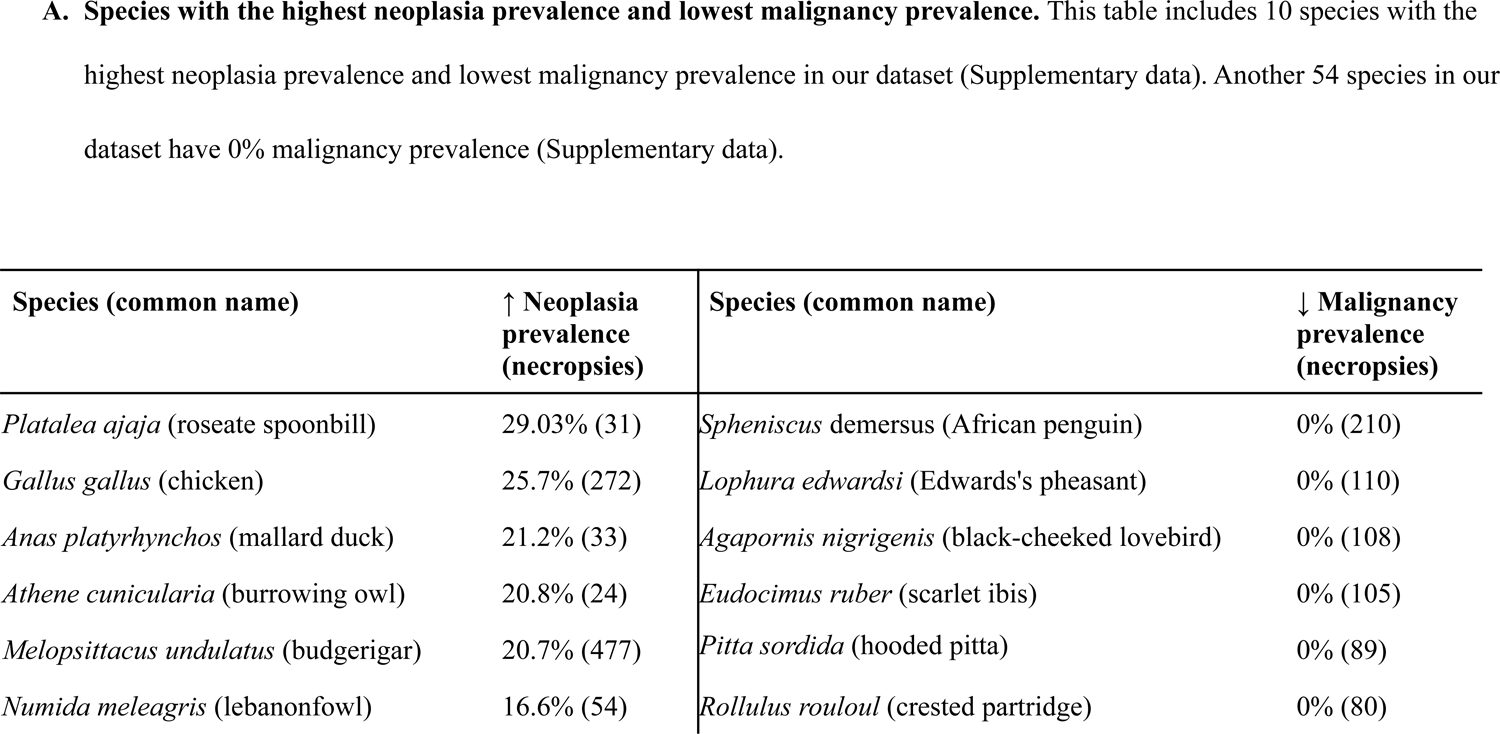
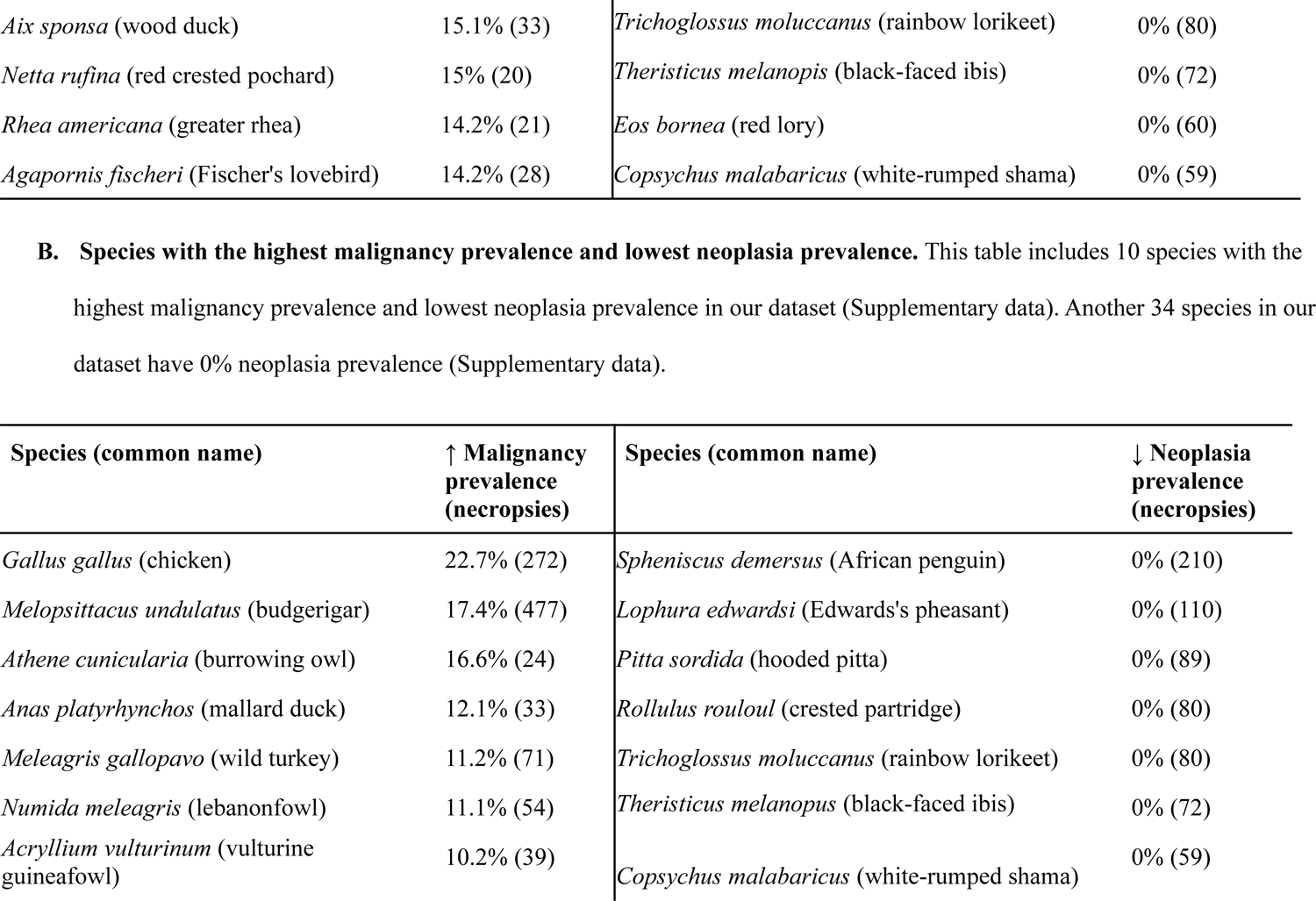
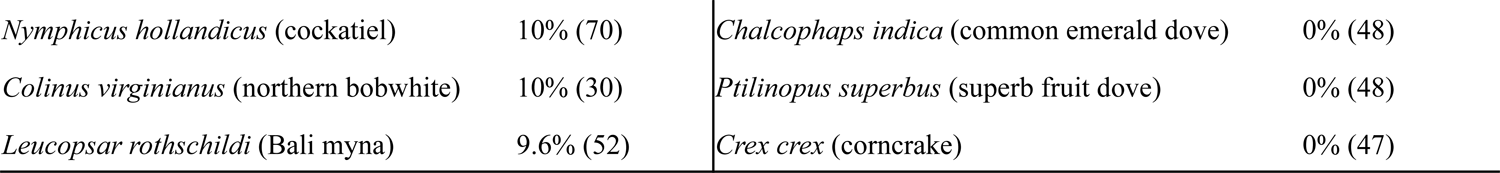
Species (A, B) with the highest and lowest malignancy prevalence and neoplasia prevalence.

We excluded all infant data from our dataset because: (1) the low prevalence of age-related diseases, such as cancer, in infants would likely bias the neoplasia prevalence data towards lower values and (2) cancers in infants are medically different than adult cancers^41^. We defined infancy as a record’s age that is smaller or equal to that species’ age of infancy (or the average of male and female maturity). In cases of no records of infancy age, the record was considered an infant if it contained any of the following words: infant, juvenile, immature, adolescent, hatchling, subadult, neonate, newborn, offspring, fledgling. We performed correlations between clutch size and neoplasia or cancer prevalence with and without removing domesticated and semi-domesticated species^42–51^ (Supplementary data). When comparing female and male malignancy prevalence and neoplasia prevalence, we removed all cases of reproductive cancer in order to minimize any effects of controlled reproduction in managed environments on our results.

### Statistical analyses

We performed all statistical analyses in R version 4.0.5^52^. We prepared figures using the data visualization software ggplot2^53^ and performed analyses in dplyr^54^. We performed all phylogenetic analyses using the R packages ape, phytools, geiger, tidyverse, powerAnalysis (https://github.com/cran/powerAnalysis), and caper^55–59^ using phylogenetic generalized least squares (PGLS) regressions to take into account the phylogenetic non-independence among species^60^ and weighting analyses by 1/(square root of the number of necropsies per species) following Revell^57^. We obtained avian phylogenetic trees from NCBI creator (https://www.ncbi.nlm.nih.gov/Taxonomy/CommonTree/wwwcmt.cgi). We performed Shapiro’s test^61^ to check for normality of the life history data and Grubbs’ & Rosner’s tests to identify and remove significant outliers. Based on the “transformTukey” function (“rcompanion” R package), which suggests a power transformation that makes values as normally distributed as possible, we log_10_-transformed the adult body mass data, log_10_-transformed the adult mass · longevity data, transformed the longevity data to the power of 0.425, and transformed clutch size (–1・clutch size–0.125).

We measured sexual differences in all seven biometric variables [plumage brightness, plumage hue, mass (g), and tail size (g)] as the natural log of the male biometric variable divided by the natural log of the female biometric variable. We also compared male malignancy prevalence or neoplasia prevalence versus female malignancy prevalence or neoplasia prevalence. The denominators in the case of the male malignancy prevalence or neoplasia prevalence are the total number of necropsied males, whereas the denominators in the case of the female malignancy prevalence or neoplasia prevalence are the total number of necropsied females. The distribution of the sex differences in cancer (i.e.,“female malignancy prevalence minus male malignancy prevalence”, “female neoplasia prevalence minus male neoplasia prevalence”) did not follow a normal distribution and had significant outliers. Therefore, we compared malignancy prevalence and neoplasia prevalence between males and females using the non-parametric paired-samples sign test. We tested whether the *P*-values passed the False Discovery Rate (FDR) correction in each of these 26 analyses (Table 2).

**Table 2.** Summary statistics. We present the summary statistics of phylogenetic regressions (PGLS) between neoplasia and malignancy prevalence and life history variables, except for the comparison of neoplasia and malignancy prevalence in females and males for which we present the summary statistics of paired-samples sign tests. The number of species analyzed is different in the majority of analyses. This is due to the fact that not all life history variables are available for every species in the literature. In the 1st P-value column we report the *P*-value of the first variable (i.e., variable A in the multivariate analysis), and in the 2nd *P*-value column we report the *P*-value of variable B. We highlight the *P-*values that passed the False Discovery Rate (FDR) correction with an asterisk (*). In the F-statistics column we report the F-statistics of variable A, and in the “Type of Association” column we report the positive (+) or negative (–) correlation between the variable A and the prevalence of neoplasia or malignancy. High lambda values show that the associations are mainly explained by common ancestry. ^†^ indicates that the R² value was not available.

| Independent variable(s) | Figure | Dependent variable | $R^2$ | F-statistic and degrees of freedom (DF) | Lambda | Type of association | *P*-value of variable A | *P*-value of variable B |
| --- | --- | --- | --- | --- | --- | --- | --- | --- |
| $\log_{10}$ adult mass | 2A | Neoplasia prevalence | 0.91 | 3.19 on 1 and 98 DF | 0.00006 | + | 0.07 | NA<sup>†</sup> |
|  | 2B | Malignancy prevalence | 0.91 | 1.09 on 1 and 98 DF | 0.22 | + | 0.29 | NA<sup>†</sup> |
| lifespan<sup>0.425</sup> | 3A | Neoplasia prevalence | 0.95 | 0.04 on 1 and 57 DF | 0.00006 | – | 0.82 | NA<sup>†</sup> |
|  | 3B | Malignancy prevalence | 0.95 | 0.15 on 1 and 57 DF | 0.00006 | – | 0.69 | NA<sup>†</sup> |
| incubation length | 4A | Neoplasia prevalence | 0.93 | 0.47 on 1 and 32 DF | 0.00006 | + | 0.49 | NA<sup>†</sup> |
|  | 4B | Malignancy prevalence | 0.93 | 2.73 on 1 and 32 DF | 0.00006 | + | 0.10 | NA<sup>†</sup> |
| –1 * clutch size<sup>–0.125</sup> | 5A | Neoplasia prevalence | 0.99 | 8.31 on 1 and 49 DF | 0.00006 | + | 0.005* | NA<sup>†</sup> |
|  | 5B | Malignancy prevalence | 0.99 | 10.80 on 1 and 49 DF | 0.01 | + | 0.0019* | NA<sup>†</sup> |
| –1 * clutch size<sup>–0.125</sup> + log<sub>10</sub> adult mass |  | Neoplasia prevalence | 0.17 | 8.38 on 1 and 48 DF | 0.00006 | + | 0.005* | 0.19 |
|  |  | Malignancy prevalence | 0.17 | 11.48 on 1 and 48 DF | 0.00006 | + | 0.0014* | 0.05 |
| –1 * clutch size<sup>–0.125</sup> | Supp. Fig. 4A | Neoplasia prevalence | 0.99 | 3.68 on 1 and 43 DF | 0.009 | + | 0.06 | NA<sup>†</sup> |
| (having excluded domesticated and semi-domesticated species) | Supp. Fig. 4B | Malignancy prevalence | 0.99 | 8.78 on 1 and 43 DF | 0.00006 | + | 0.004* | NA<sup>†</sup> |
| -1 * clutch size<sup>-0.125</sup> + log<sub>10</sub> adult mass (having excluded domesticated and semi-domesticated species) |  | Neoplasia prevalence | 0.08 | 3.91 on 1 and 42 DF | 0.00006 | + | 0.05 | 0.35 |
|  |  | Malignancy prevalence | 0.1 | 8.9 on 1 and 42 DF | 0.00006 | + | 0.004* | 0.21 |
| degree of dimorphism in brightness | 6A | Neoplasia prevalence | 0.75 | 1.00 on 1 and 16 DF | 0.11 | + | 0.33 | NA<sup>†</sup> |
|  | 6B | Malignancy prevalence | 0.73 | 0.09 on 1 and 16 DF | 0.33 | + | 0.76 | NA<sup>†</sup> |
| degree of dimorphism in hue | 6C | Neoplasia prevalence | 0.44 | 0.09 on 1 and 22 DF | 0.10 | - | 0.76 | NA<sup>†</sup> |
|  | 6D | Malignancy prevalence | 0.41 | 0.15 on 1 and 22 DF | 0.35 | - | 0.69 | NA<sup>†</sup> |
| degree of dimorphism in mass | 6E | Neoplasia prevalence | 1 | 1.14 on 1 and 45 DF | 0.24 | – | 0.28 | NA<sup>†</sup> |
|  | 6F | Malignancy prevalence | 1 | 0.10 on 1 and 45 DF | 0.32 | – | 0.74 | NA<sup>†</sup> |
| degree of dimorphism in tail size | 6G | Neoplasia prevalence | 1 | 0.03 on 1 and 32 DF | 0.40 | – | 0.84 | NA<sup>†</sup> |
|  | 6H | Malignancy prevalence | 1 | 0.11 on 1 and 32 DF | 0.60 | + | 0.73 | NA<sup>†</sup> |
| sex | 7A | Neoplasia prevalence | 97.1% CI = -0.05 - 0% |  |  |  | 0.16 | NA<sup>†</sup> |
|  | 7B | Malignancy prevalence | 97.1% CI = 0 - 0.01% |  |  |  | 0.66 | NA<sup>†</sup> |
| log<sub>10</sub> (adult mass * lifespan) | Supp. Fig. 1A | Neoplasia prevalence | 0.97 | 0.06 on 1 and 55 DF | 0.00006 | + | 0.79 | NA<sup>†</sup> |
|  | Supp. Fig. 1B | Malignancy prevalence | 0.97 | 0.001 on 1 and 55 DF | 0.00006 | – | 0.97 | NA<sup>†</sup> |

## Results

The range of neoplasia prevalence among the examined 108 bird species varied from 0% to 29%, with a mean of 4.4%, whereas malignancy prevalence among these species varied from 0% to 17.4%, with a mean of 2.3% (Table 1; supplementary data). Among the four avian taxonomic orders with at least 10 species per order in our dataset (Psittaciformes, Passeriformes, Columbiformes, and Anseriformes), the Anseriformes had on average the highest malignancy prevalence (mean ± SD: 2.84% ± 2.81%), whereas the Columbiformes had on average the lowest malignancy prevalence (mean ± SD: 1.12% ± 1.84%) (Supplementary data). We found no significant correlation between neoplasia or malignancy prevalence: and (1) adult body mass across 100 bird species and 5042 necropsies (Fig. 2A; Fig. 2B; Table 2); nor (2) adult mass times lifespan across 57 bird species and 3464 necropsies (Supp. Fig. 1A; Supp. Fig. 1B: Table 2). Neoplasia and malignancy prevalence were not higher in longer-lived birds (Fig. 3A; Fig. 3B: Table 2; 59 species and 3593 necropsies), and deaths with a necropsy diagnosis of cancer were not skewed towards old age across 1287 individuals from 51 species (Supp. Fig. 3).

**Figure 1.**
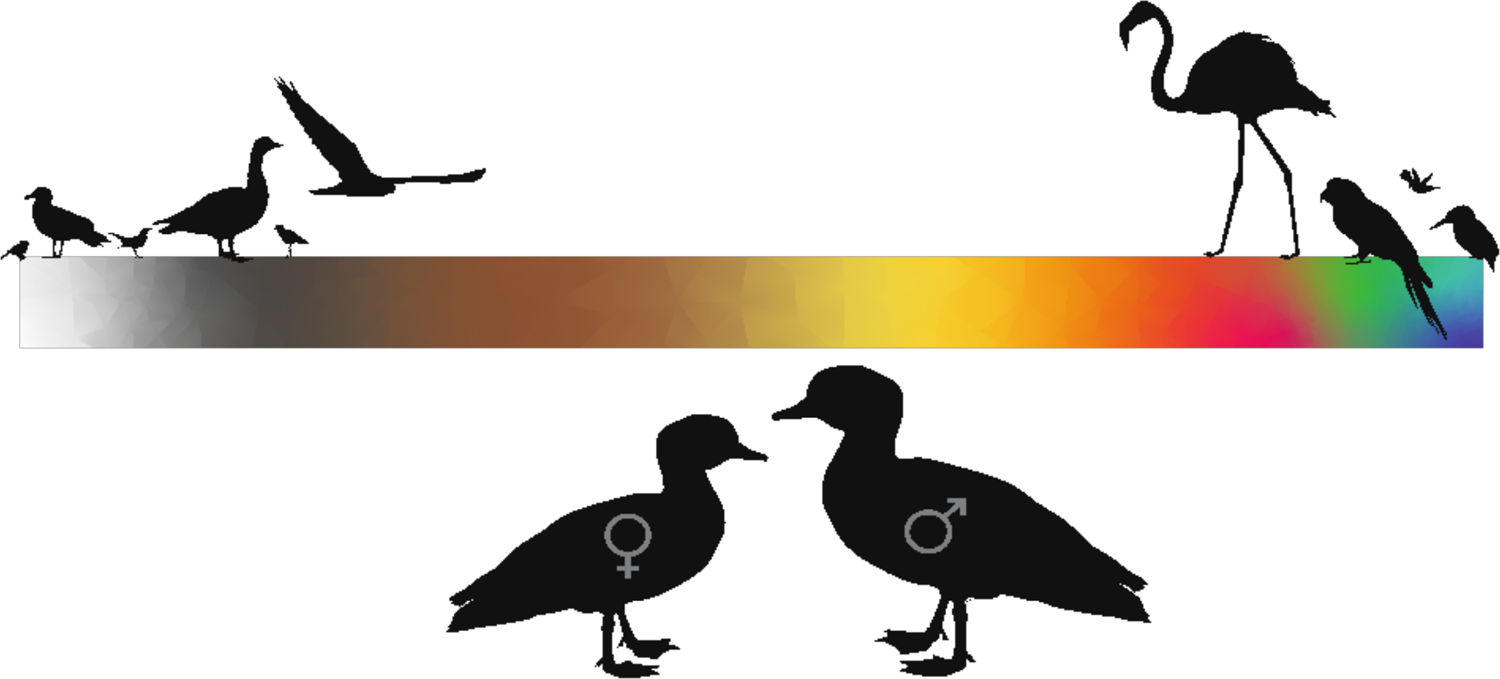
Sexual dimorphism in birds. Birds display a wide range of sexual dimorphism in size and plumage color.

**Figure 2.**
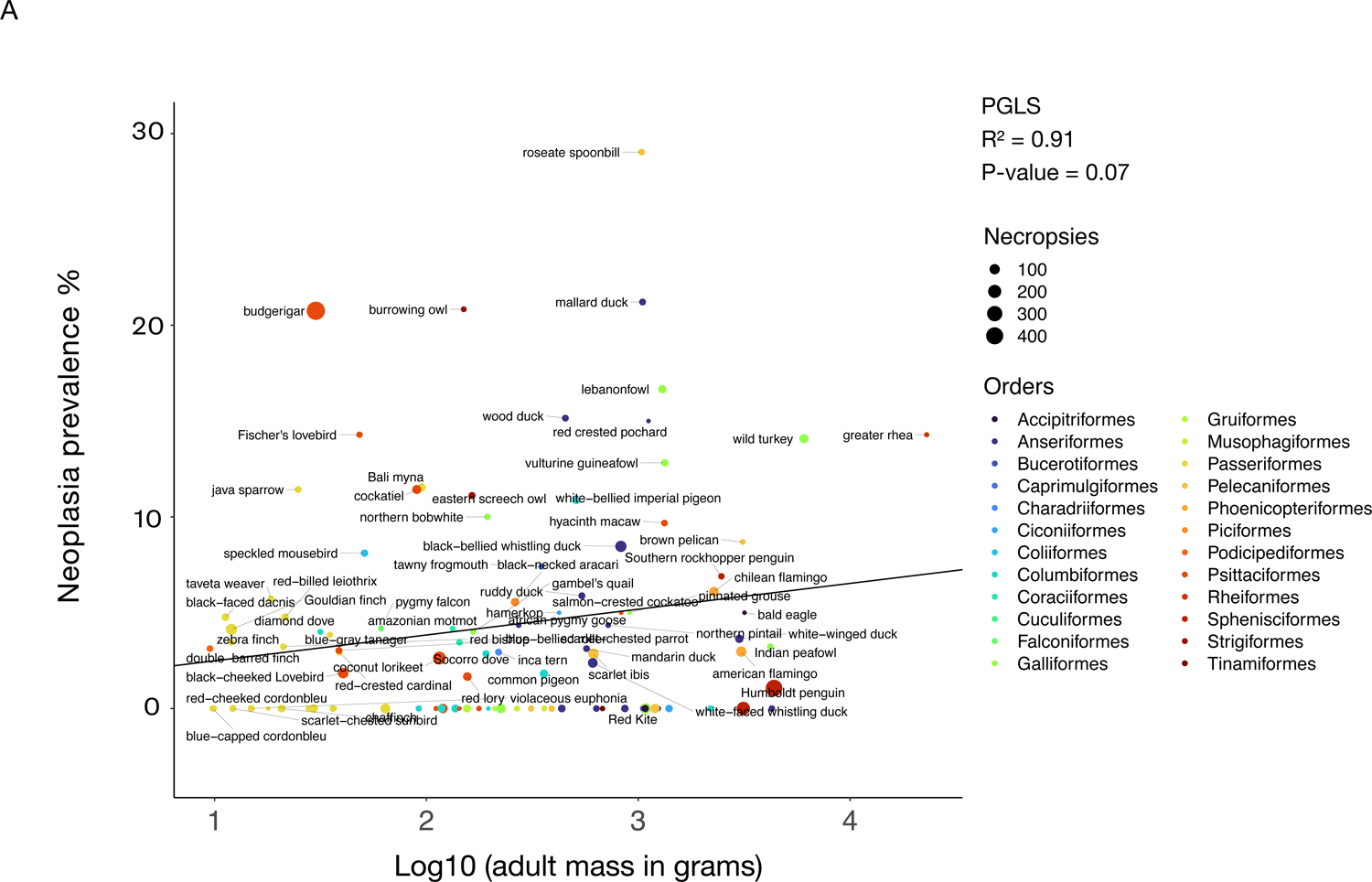
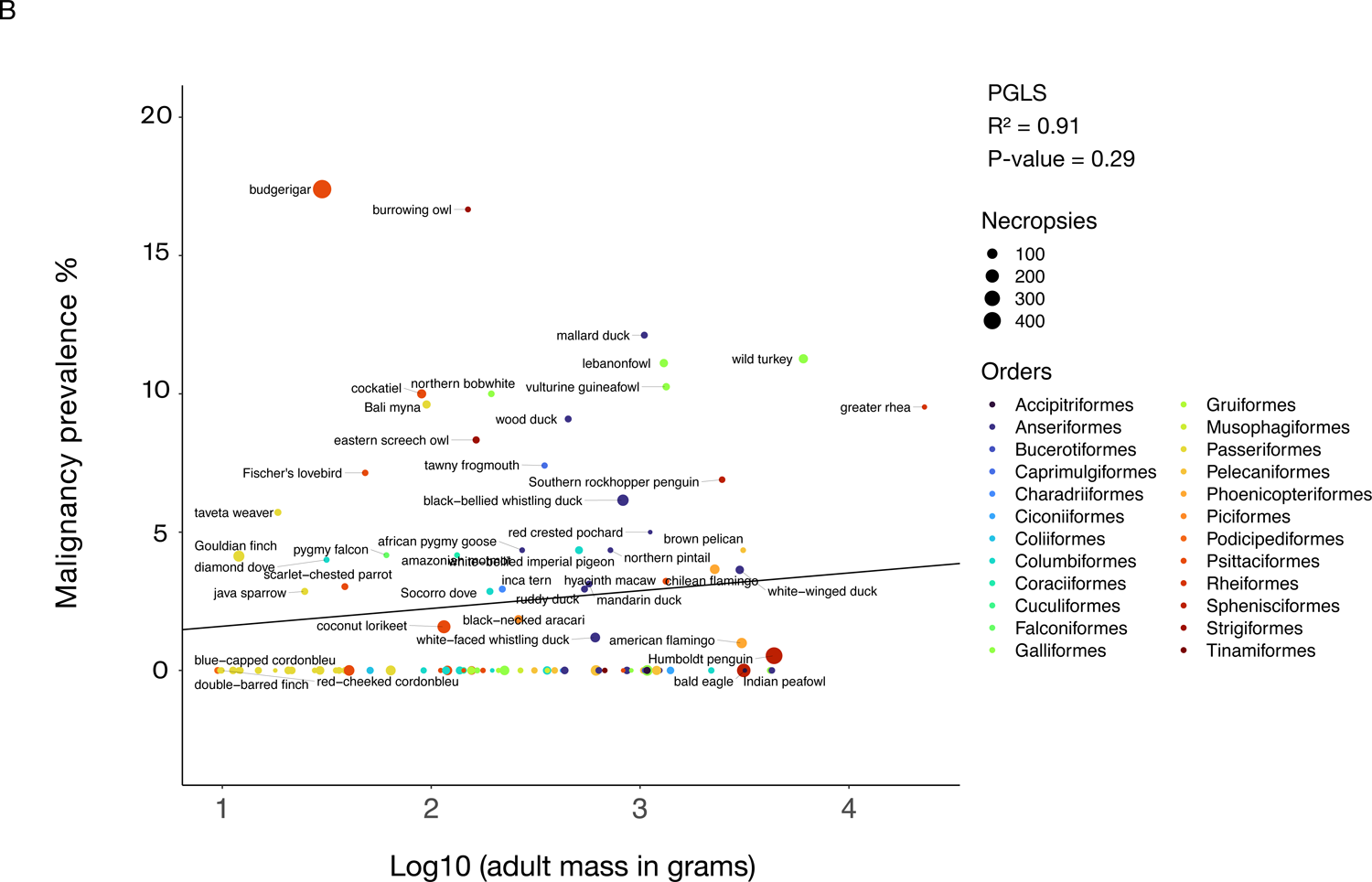
Larger body mass is not correlated with neoplasia prevalence (A) or malignancy prevalence (B) across 100 bird species. Dot size indicates the number of necropsies per species. Colors show the taxonomic order of each species, and black lines show the phylogenetically-controlled linear regression of the logarithm of adult mass versus malignancy prevalence or neoplasia prevalence.

**Figure 3.**
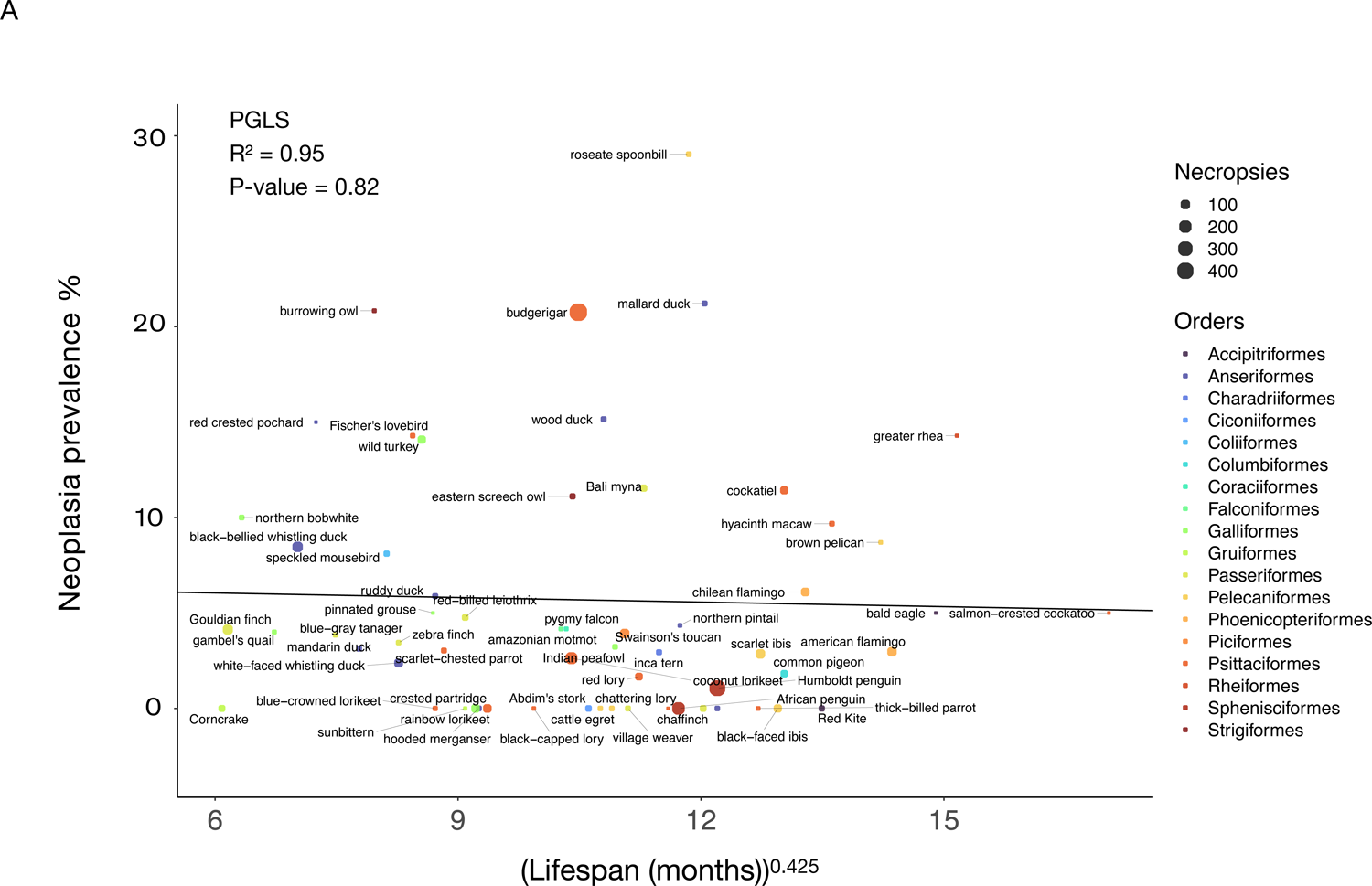
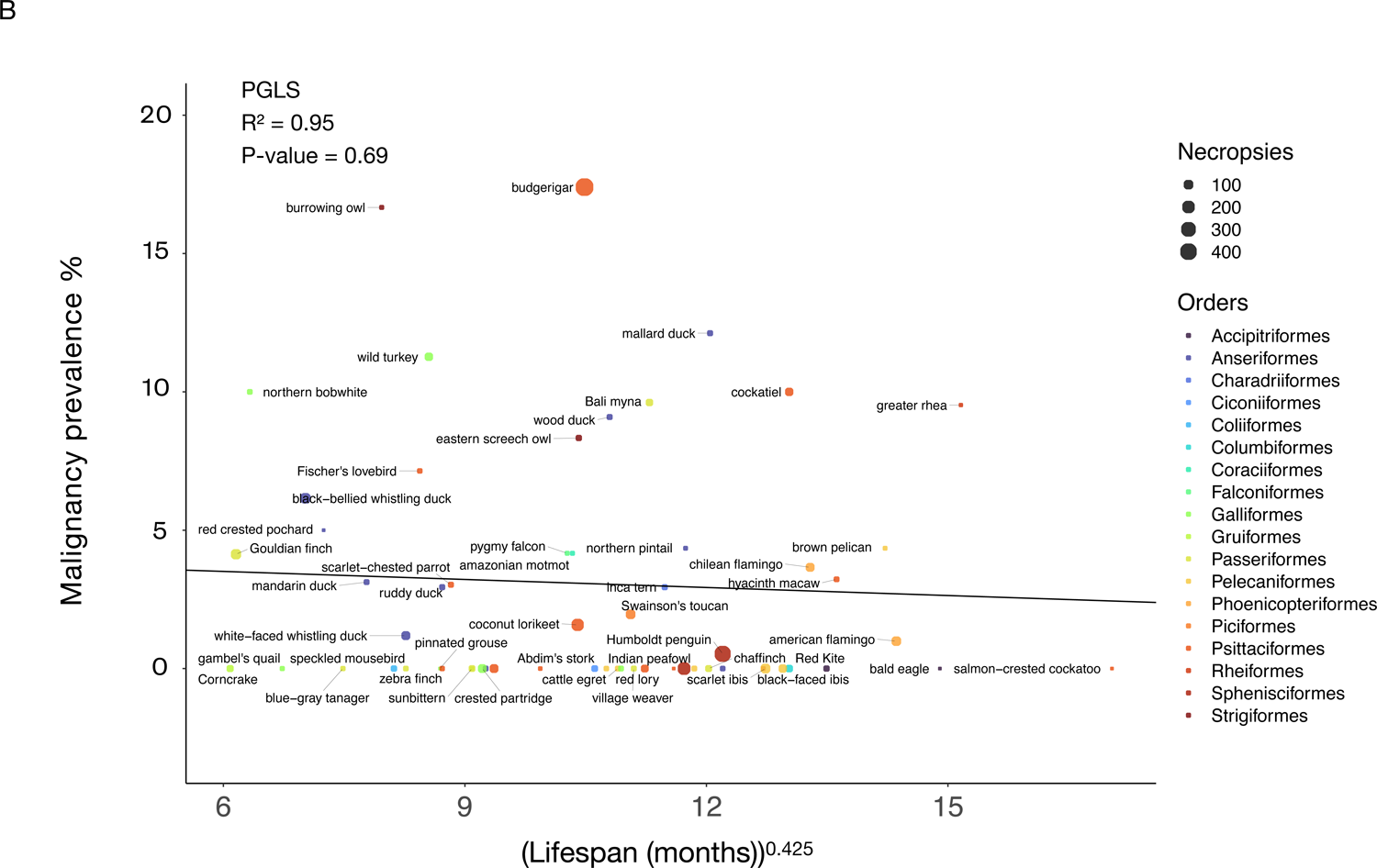
Longer lifespan is not correlated with neoplasia prevalence (A) or malignancy prevalence (B) across 59 bird species. Dot size indicates the number of necropsies per species. Colors show the taxonomic order of each species. Black lines show the phylogenetically-controlled linear regression of the normalized values of species lifespan versus malignancy prevalence or neoplasia prevalence.

We found that length of incubation was not significantly correlated with neoplasia or malignancy prevalence (Fig. 4A; Fig. 4B; Table 2; 34 species and 1806 necropsies). However, species with larger clutch sizes had significantly higher neoplasia and malignancy prevalence even after applying FDR corrections for multiple testing (*P-*value = 0.005, R² = 0.99; and *P-*value = 0.0019, R² = 0.99, respectively; Fig. 5; 51 species and 2119 necropsies), and after controlling for species body mass (*P-*value = 0.005, R² = 0.17; and *P-*value = 0.0014, R² = 0.17, respectively; Table 2). The positive correlation between clutch size and malignancy prevalence, but not neoplasia prevalence, remained significant after removing domesticated and semi-domesticated species (*P-*value = 0.004, R² = 0.99; Supp. Fig. 4B; 45 species and 1839 necropsies) and controlling for body mass (*P-*value = 0.004, R² = 0.1; Table 2; 45 species). We found no significant associations between neoplasia or malignancy prevalence and several sexually dimorphic and dichromatic traits (Fig. 6; Table 2). Also, neoplasia and malignancy prevalence were not significantly different between males and females across 31 species (Fig. 7; Supp. Fig. 2; Table 2).

**Figure 4.**
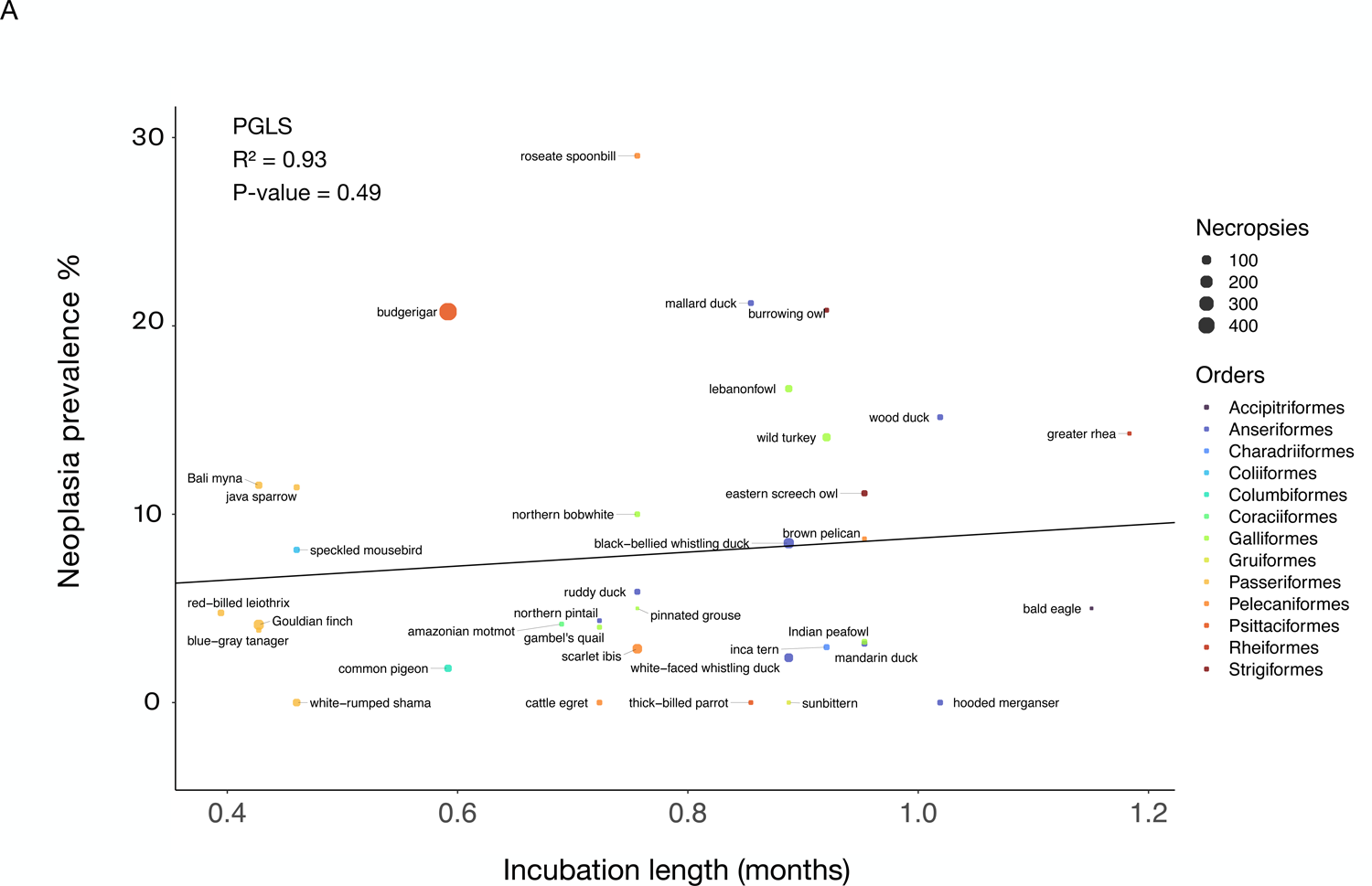
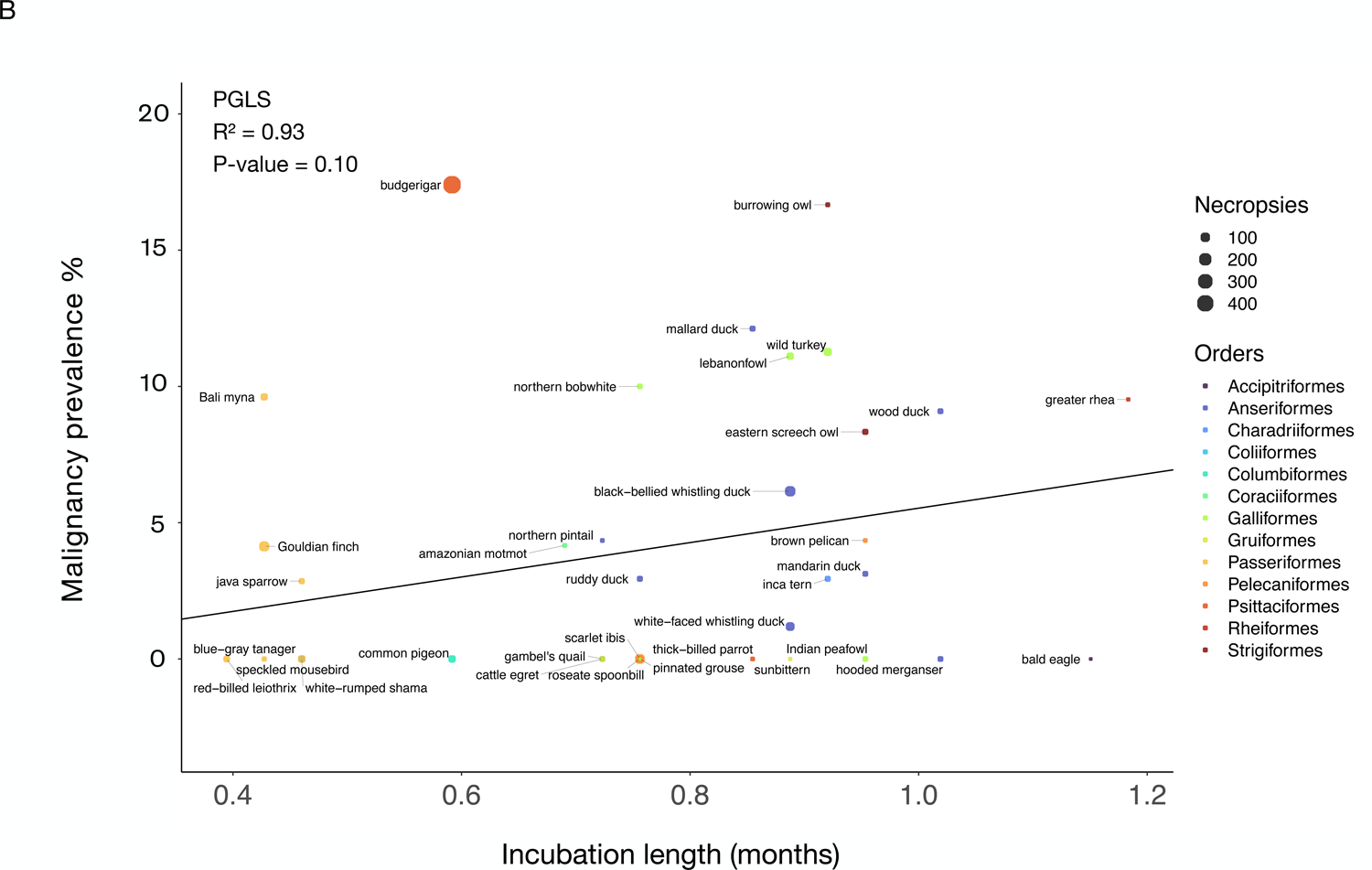
Incubation length is not correlated with neoplasia prevalence (A) or malignancy prevalence (B) when controlling for body mass across 34 bird species. Different colors indicate the order in which each species belongs and the size of the dot indicates the number of necropsies per species. Black lines show the phylogenetically-controlled linear regression of incubation length versus malignancy prevalence or neoplasia prevalence.

**Figure 5.**
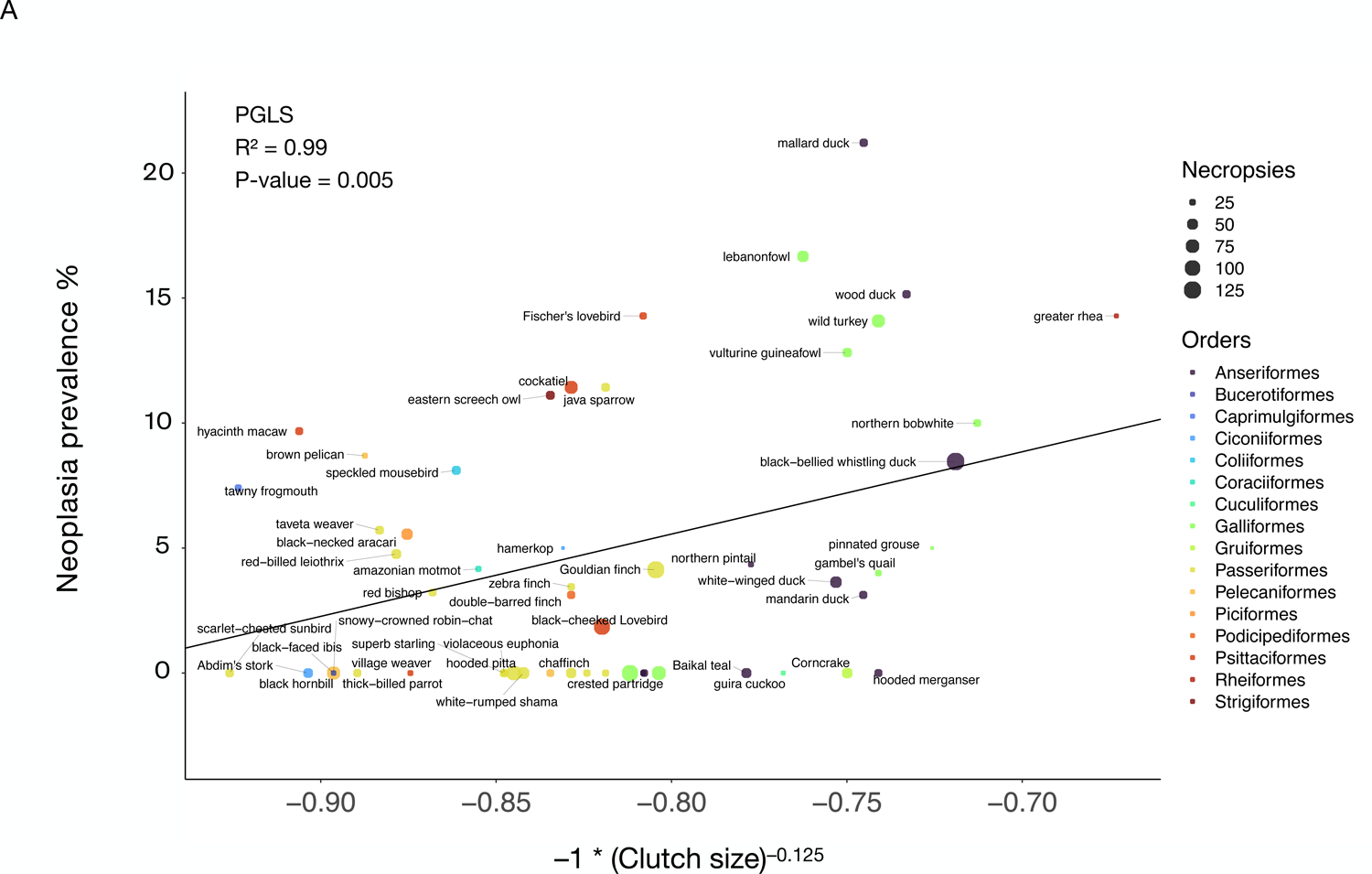
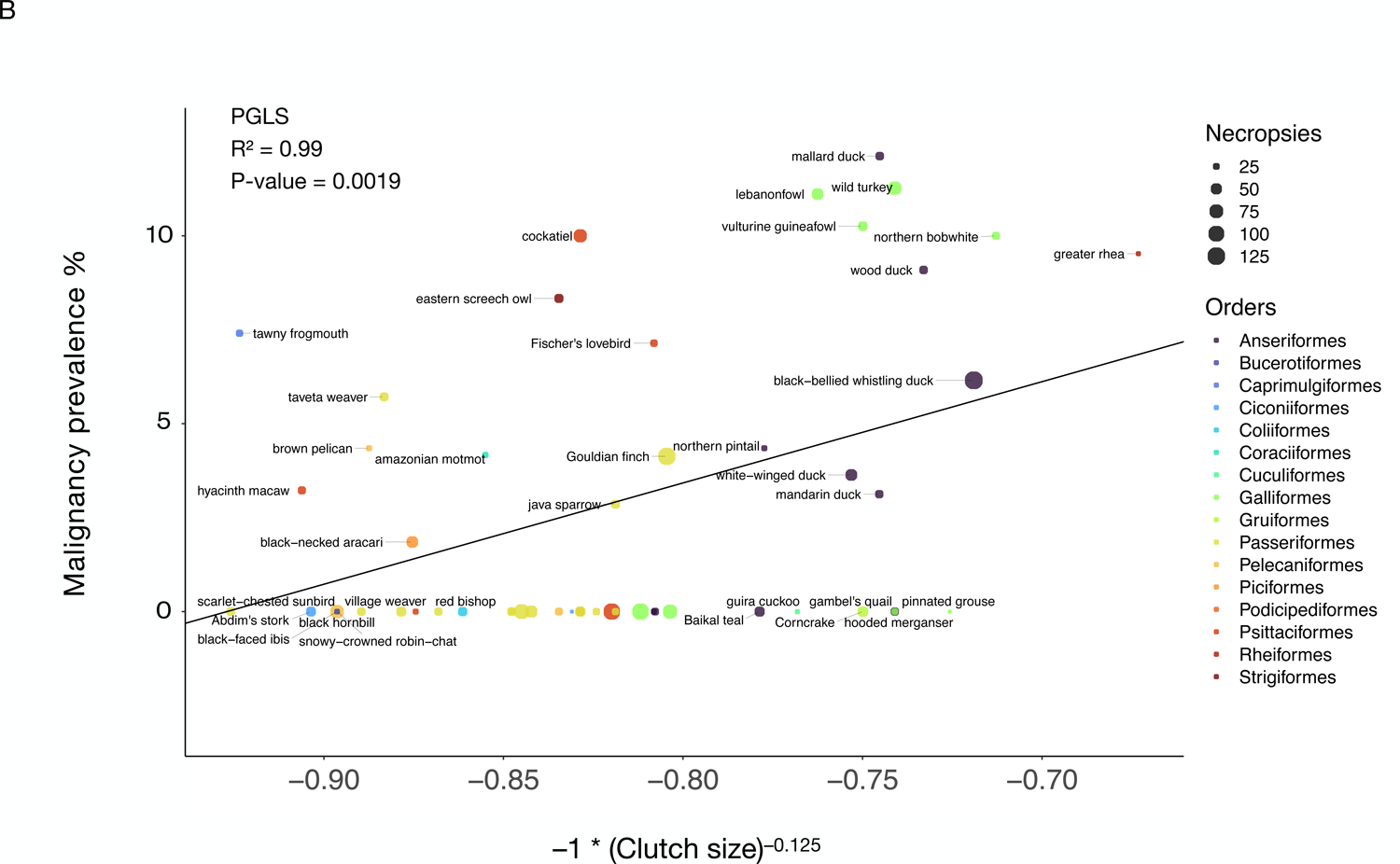
Larger clutch size is correlated with neoplasia prevalence (A) and malignancy prevalence (B) across 51 bird species. After controlling for species body mass, the positive correlation between clutch size and neoplasia prevalence (*P*-value = 0.005; Table 2) and malignancy prevalence (*P*-value = 0.0014; Table 2) remains significant. Dot size indicates the number of necropsies per species. Colors show the taxonomic order of each species. Black lines show the phylogenetically-controlled linear regression of the normalized values of clutch size versus malignancy prevalence or neoplasia prevalence.

**Figure 6.**
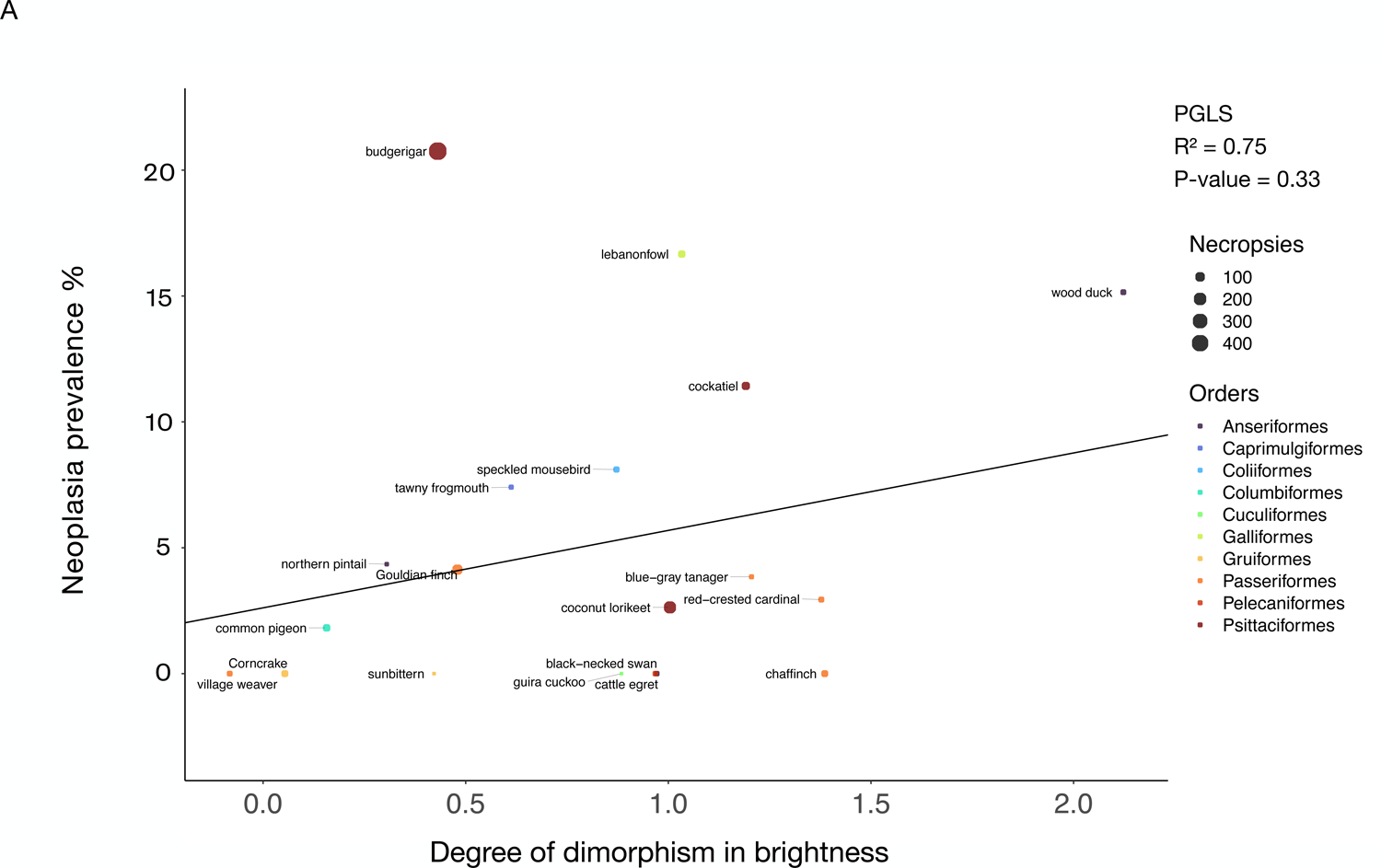
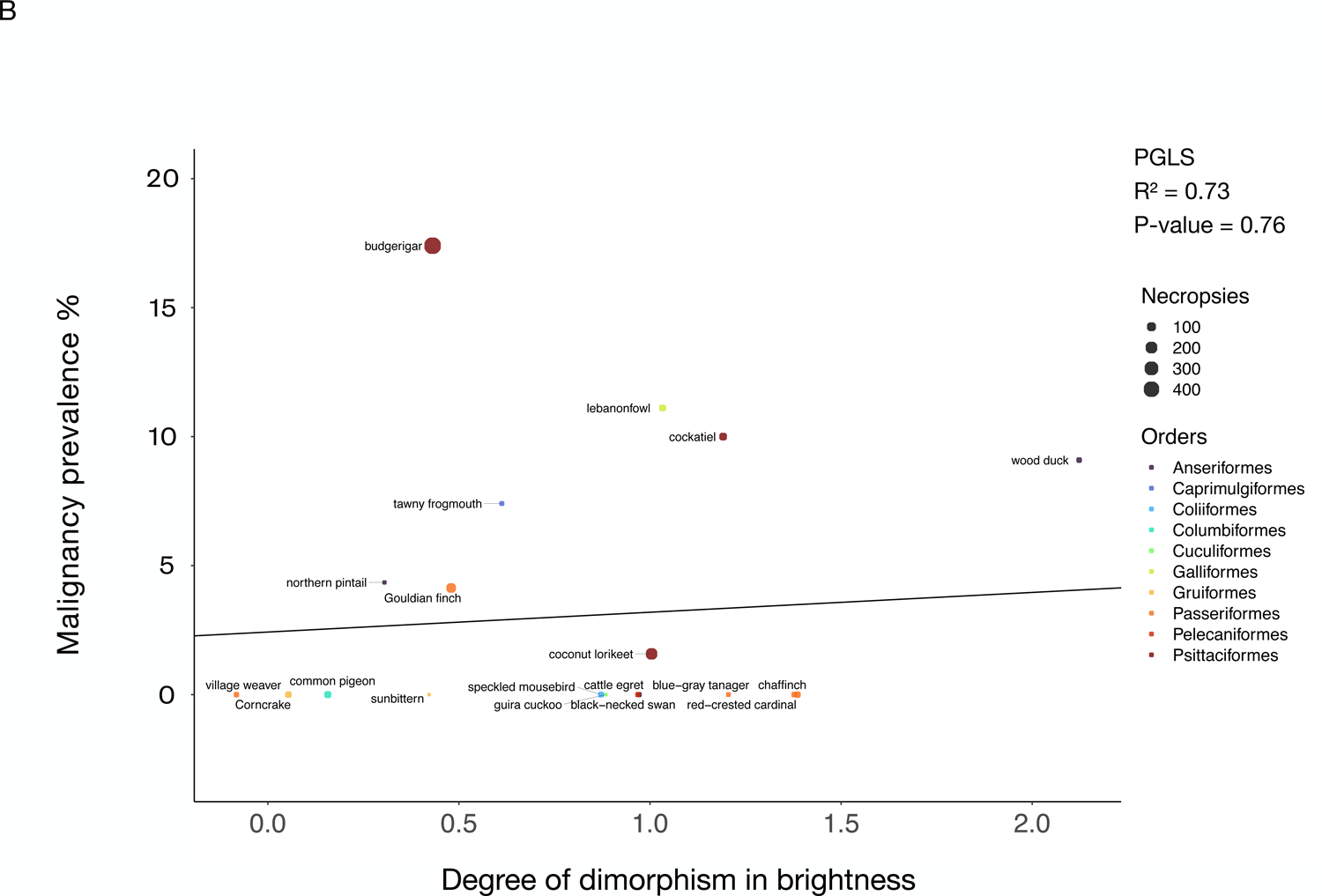
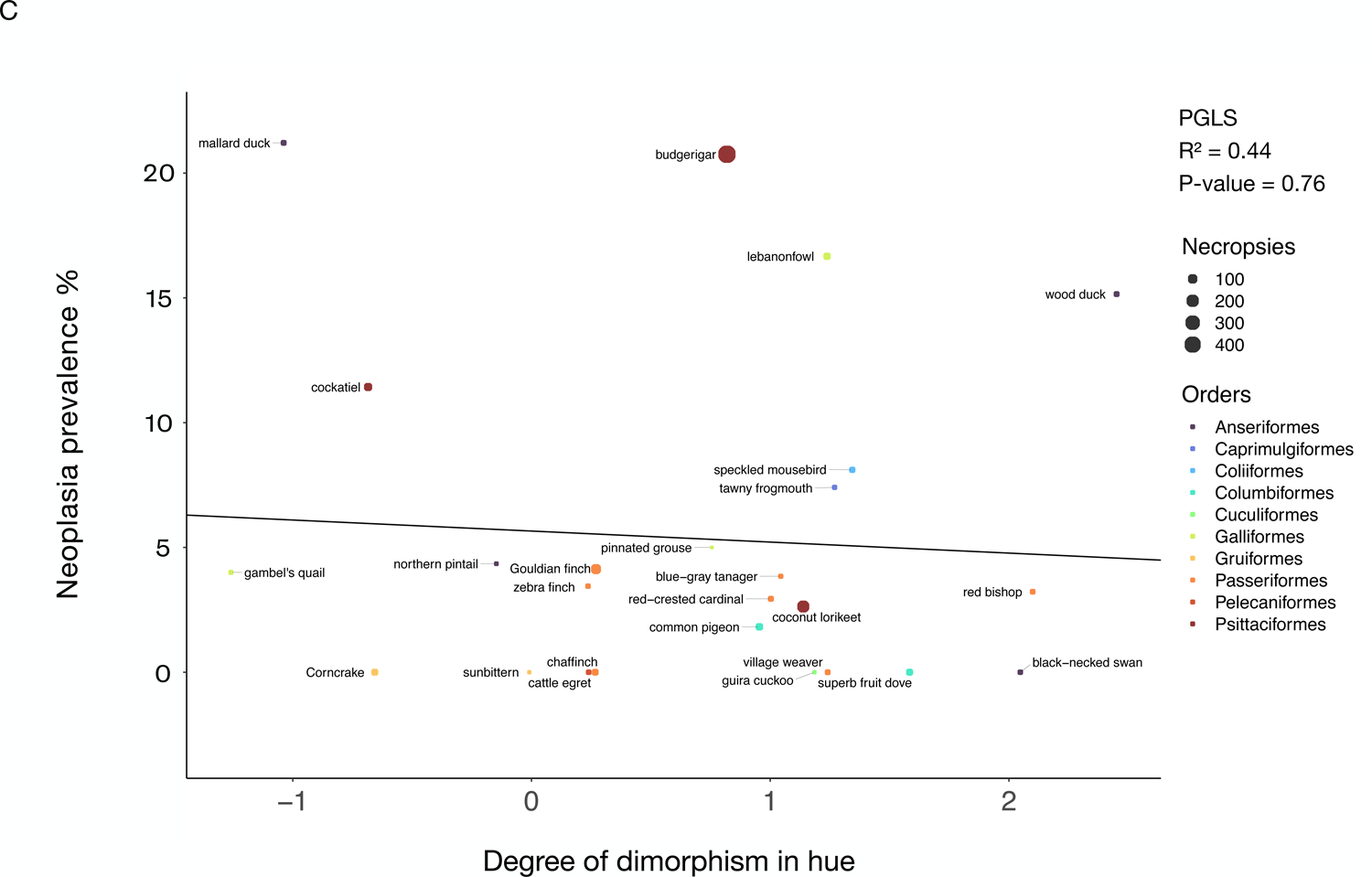
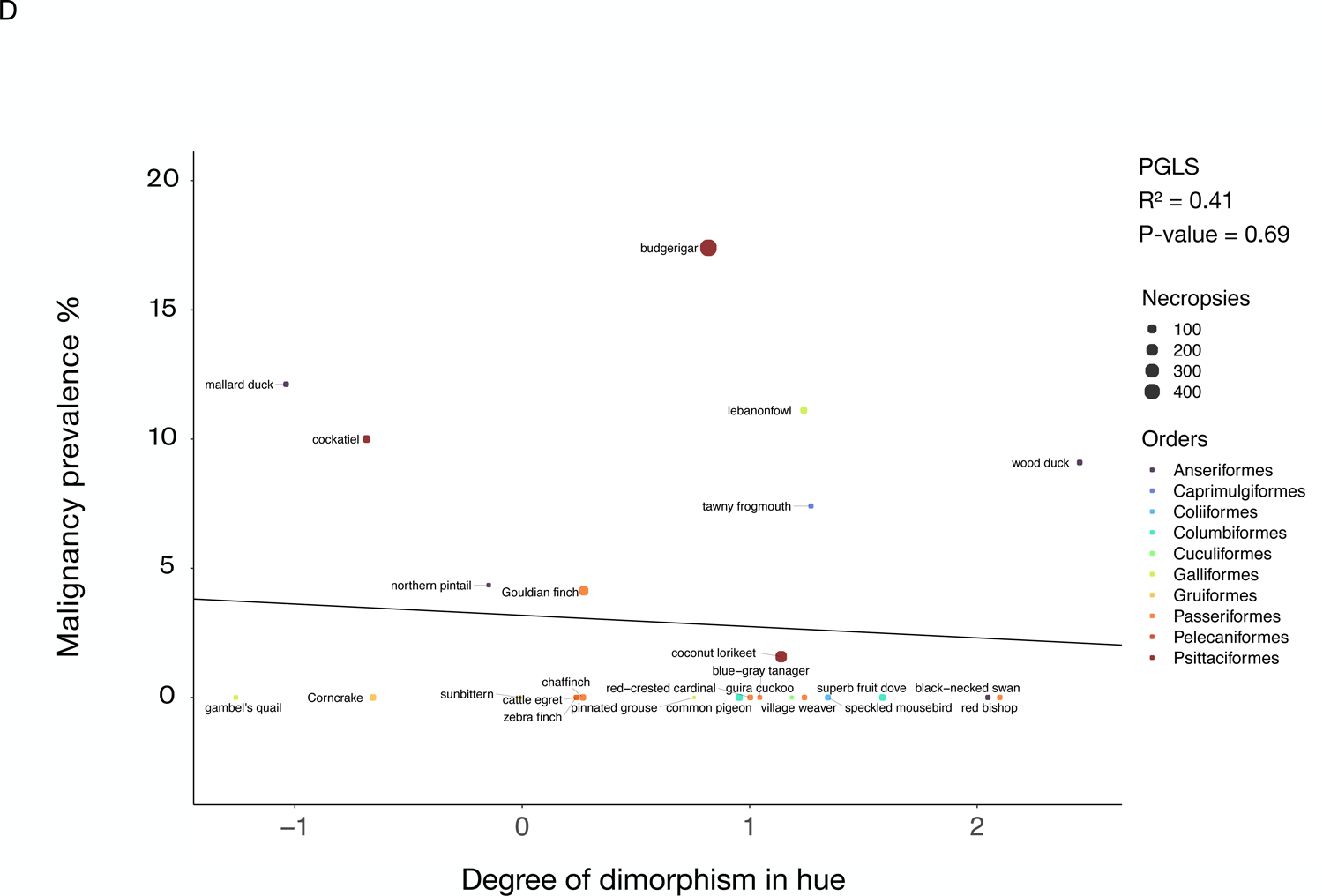
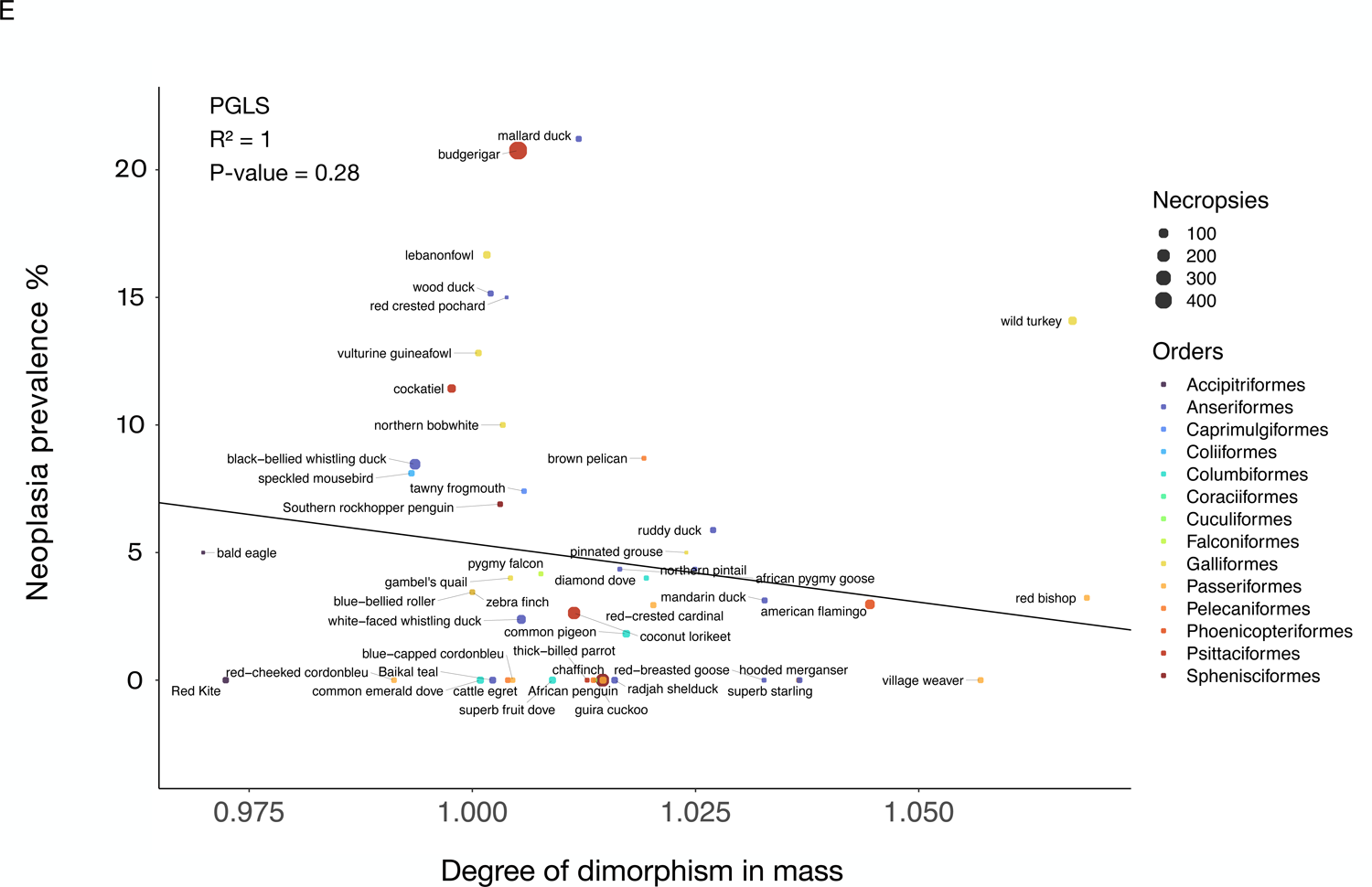
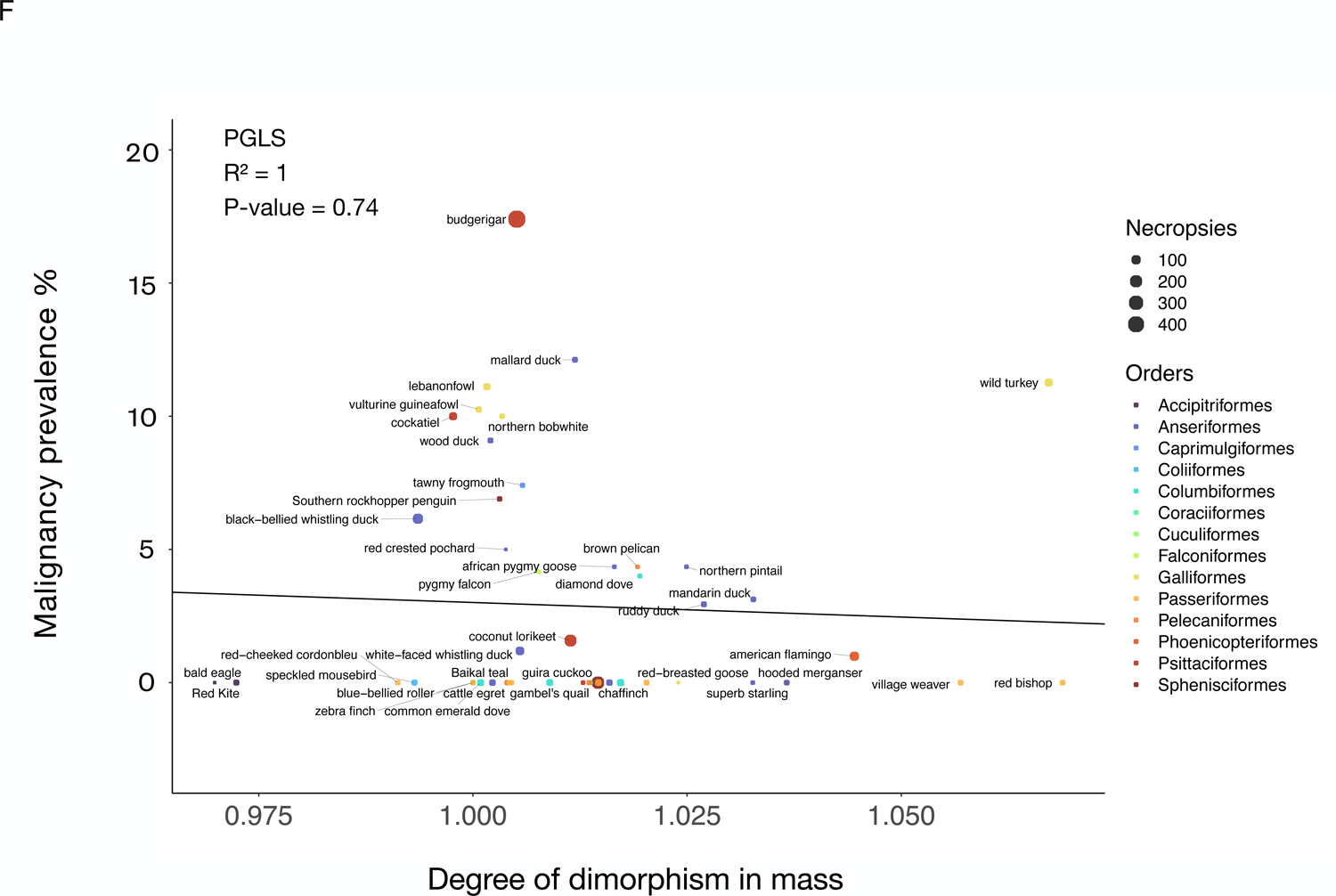
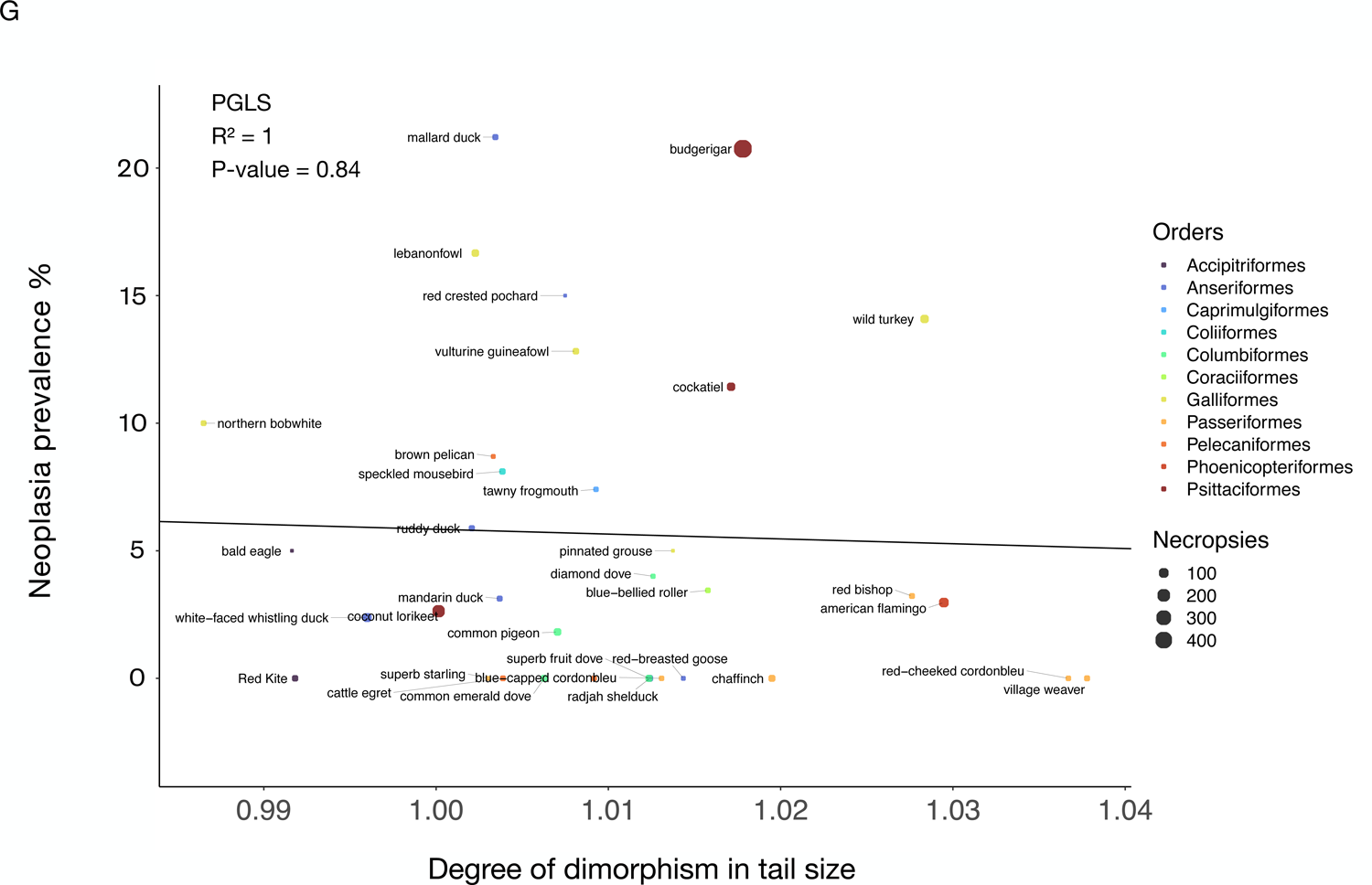
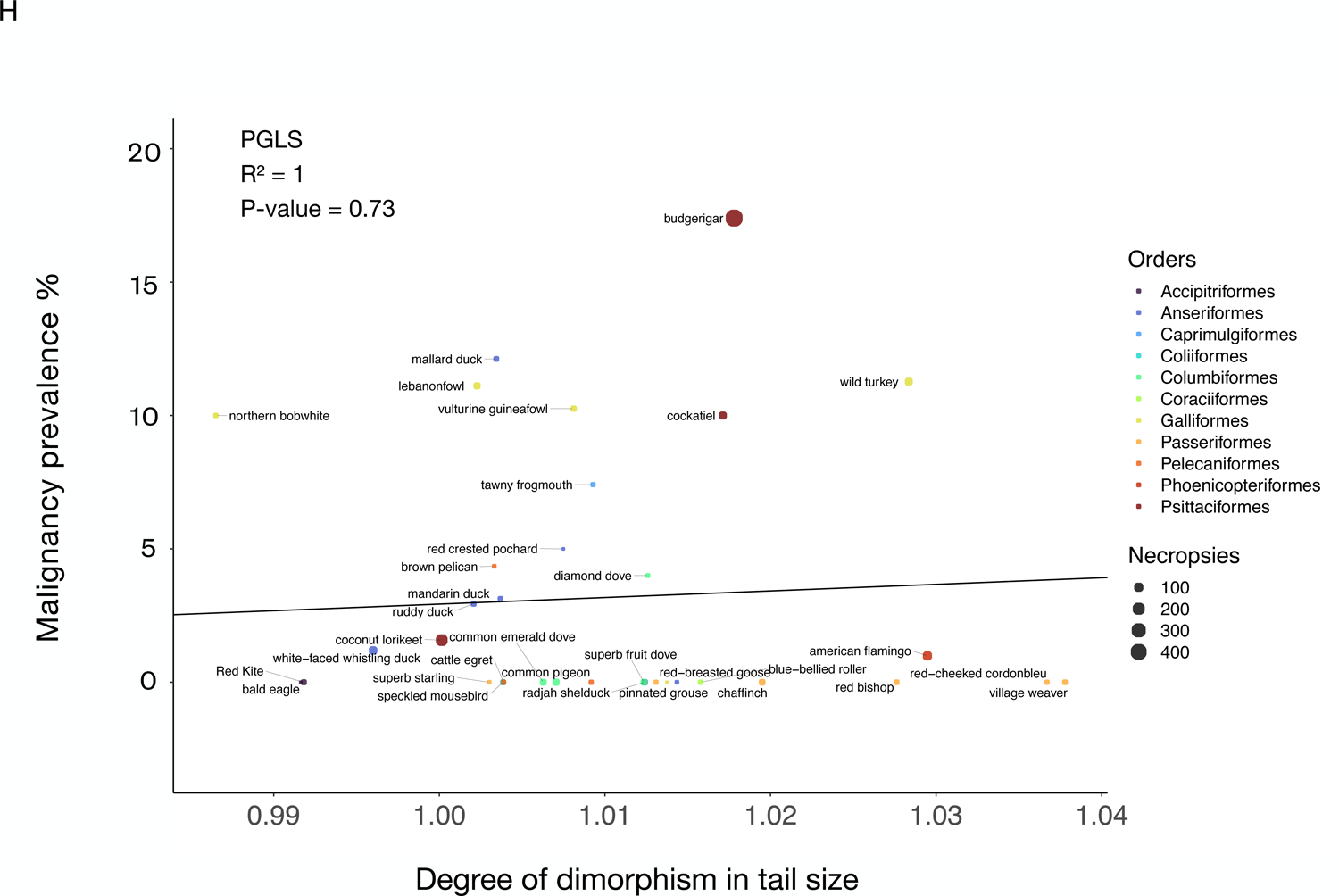
Sexual dimorphic traits are not correlated with neoplasia or malignancy prevalence in birds. The degree of dimorphism in brightness is not correlated with neoplasia prevalence (A) or malignancy prevalence (B) across 18 species of birds. The degree of dimorphism in hue is not correlated with neoplasia prevalence (C) or malignancy prevalence (D) across 24 species of birds. The degree of dimorphism in mass is not correlated with neoplasia prevalence (E) or malignancy prevalence (F) across 47 species of birds. The degree of dimorphism in tail size is not correlated with neoplasia prevalence (G) or malignancy prevalence (H) across 34 species of birds. A positive score on the x-axis indicates that the species has a relatively higher score in that trait in males than females, whereas a negative score on the x-axis shows that the species has a relatively higher score in that trait in females than males. Black lines show the phylogenetically-controlled linear regression of the degree of dimorphism in the trait versus neoplasia prevalence or malignancy prevalence. Different colors indicate the order in which each species belongs and the size of the dot indicates the total number of necropsies per species.

**Figure 7.**
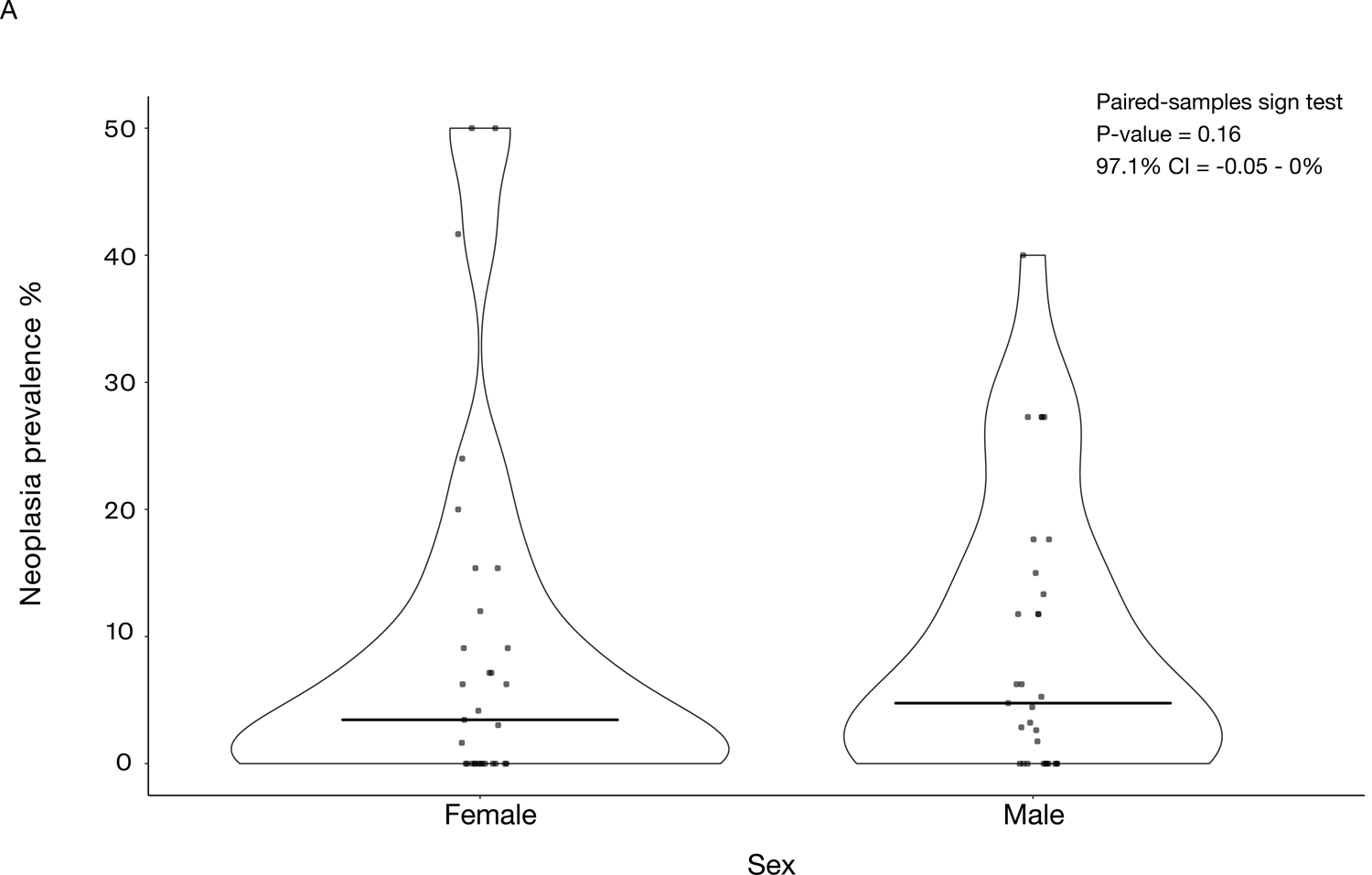
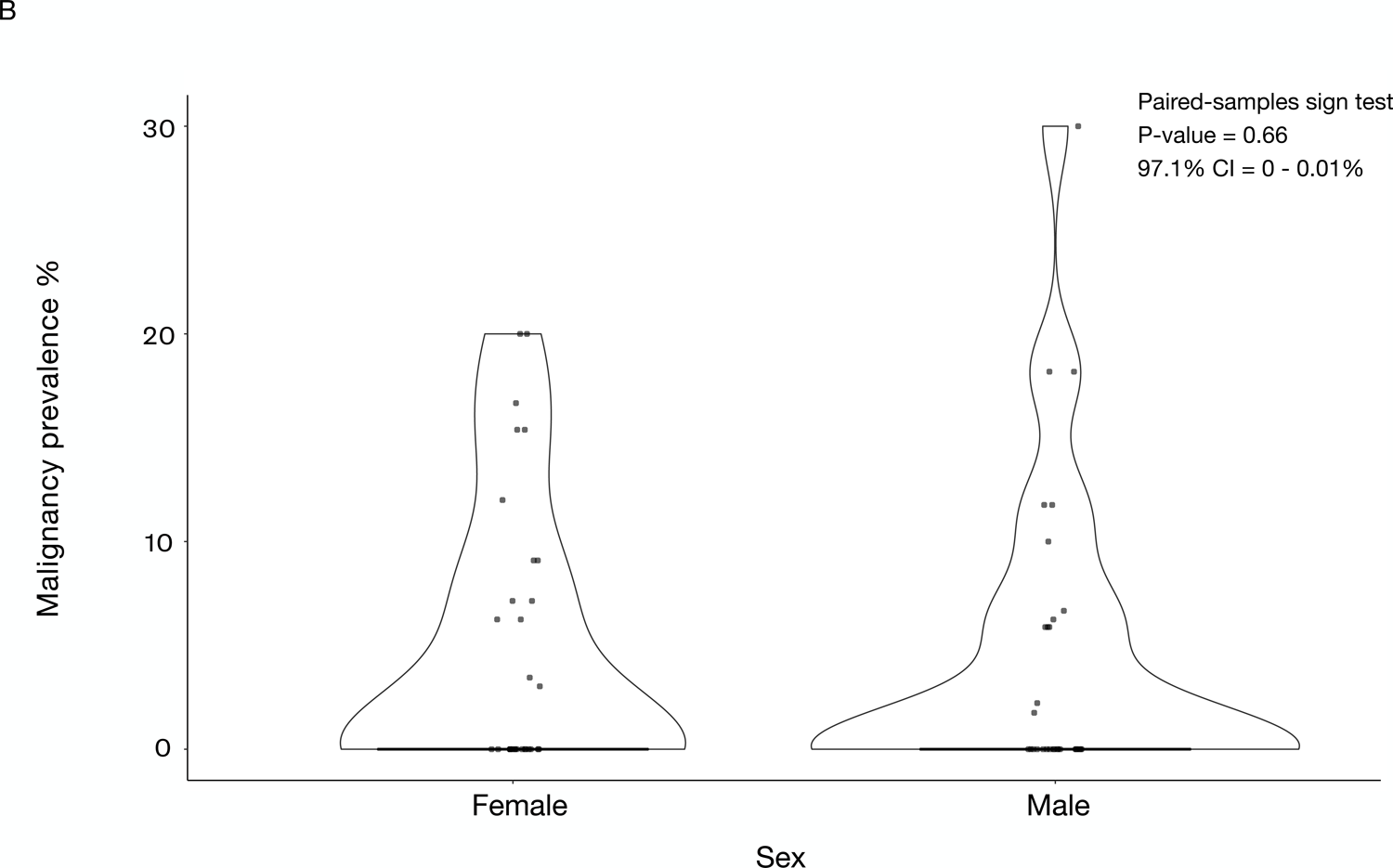
Neoplasia (A) and malignancy prevalence (B) are not significantly different between females and males across 31 bird species. Horizontal bars show the median neoplasia (A) or malignancy prevalence (B). We added minimal jitter for better visualization of individual data points.

## Discussion

We hypothesized that differences in life-history traits, including clutch size, may explain some of the variation in cancer prevalence across managed bird species. Species varied in their clutch sizes from scarlet-chested sunbirds laying on average 1.85 eggs, to greater rheas laying >10 times as many (23 eggs on average). We found that clutch size explained a statistically significant portion (17%) of the variation in cancer prevalence when controlling for log_10_ adult mass. Species with larger clutch size had higher malignancy and neoplasia prevalence, even after FDR corrections and controlling for body mass. The positive correlation between clutch size and malignancy prevalence remained significant even after removing domesticated and semi-domesticated species from the analysis. However, no other life-history trait that we measured, such as adult body mass, lifespan, incubation length, sexual size dimorphism or sexual dichromatism, explained the variance in avian cancer prevalence, nor was there a significant difference in cancer or neoplasia prevalence between male and female birds.

### Body mass and lifespan are not associated with cancer in birds managed under human care

Our observations in populations of birds managed under human care show no significant correlation between neoplasia or malignancy prevalence and adult body mass, lifespan, or adult mass times lifespan in birds, supporting Peto’s paradox^62^; however, these results are in contrast to the observation of cancer in free-living birds^19^. While there is a trend in our data for larger birds to have more cancer, this was not statistically significant (*P-*value = 0.29). The discrepancy between our study and that of Møller et al.^19^, may be due to the different number of individuals sampled per species (≥3 records per species in Møller et al.^19^ versus ≥20 necropsies per species in our study), the different species of birds analyzed (238 free-living bird species in Denmark^19^ versus 108 managed bird species from multiple institutions), or body mass mostly measured with a precision balance^19^ versus collected from the literature. In addition, birds collected by Mølller et al. were mostly killed by hunters (both human and non-human), whereas those in our study were protected from predation and thus allowed to live long enough to succumb to various diseases of old age, including cancer. Unfortunately, only six species of birds are common in Møller et al.’s^19^ and this study’s dataset, limiting our ability to compare cancer prevalence in wild versus managed birds. In general, patterns of tumor incidence or neoplasia prevalence were consistent between these free-living birds and populations managed under human care (Supplementary Table 1). Therefore, while there are many potential sources of error in the enumeration of the life-history traits and neoplasia prevalence in either wild or managed birds, it is promising that there is consistency in shared data trends across studies.

Interestingly, the roseate spoonbill, ranked 18th among the oldest species with lifespan data in our dataset, has the highest neoplasia prevalence (29.03%), but no reported malignancy (0% malignancy prevalence). We found that birds that live longer do not have significantly higher cancer prevalence than shorter-lived species, and there is not a skew in terms of more cancer deaths towards old age. This may be explained by the observation that long-lived birds have coevolved pathways that increase longevity in part through decreasing cancer rates^63, 64^. Specifically, in long-lived birds, there is an increased selective pressure for genes related to controlling cell division and tumor suppression^63^. Long-lived mammals, such as bats, have extra copies of *FBXO31* and mutations in the insulin-like growth factor 1 receptor/growth-hormone receptor related to blocking the cell cycle and responding to DNA damage^65–67^. The fact that erythrocyte telomeres of long-lived birds shorten at a slower pace than erythrocyte telomeres of shorter-lived birds^68^ may provide an additional mechanistic explanation for the lower than expected cancer prevalence in long-lived birds.

### Neoplasia and cancer prevalences are higher in species with larger clutch sizes

Our results are consistent with previous findings that larger litter size is associated with cancer prevalence in mammals^35, 69^. Many of the life-history traits described in this article, such as body mass, number of offspring produced, incubation time, and longevity, are tightly linked with each other^70–74^ (Supp. Fig. 5). No significant correlations were found between cancer prevalence and lifespan, adult mass, or incubation/gestation length in birds or mammals^35^. Larger clutch size is correlated with malignancy prevalence and neoplasia prevalence, even after corrections for multiple testing and controlling for species body mass. This discrepancy between clutch size predicting neoplasia prevalence but not the other (correlated) life-history variables may be due to the fact that we only have clutch size data on a subset of the species (51 bird species) for which we have other life-history data (e.g., 100 bird species with adult mass data). It could also be that distinct molecular pathways associated with clutch size have coevolved with increased neoplasia and malignancy prevalence.

We found that when including domesticated species in the analyses, both malignancy and neoplasia prevalence are positively correlated with clutch size, however, when excluding domesticated species, only malignancy prevalence remains positively correlated with clutch size; indicating that differential selection pressures may be acting on neoplasia versus malignancy. In some birds kept in enclosures with artificial light, we speculate that the exposure to artificial light could be one explanation for the association between neoplasia prevalence and clutch size when domesticated and semi-domesticated species are included in the analysis. Artificial light is used in poultry industries, as well as parakeet breeding, to lengthen the hours of egg laying^75, 76^, and such prolonged exposure to light of high intensity has been suggested to cause hyperplasia and neoplasia in the pituitary^76^.

### Is sexual dimorphism or dichromatism correlated with cancer prevalence in birds?

The strength of sexual selection could impose energetic constraints resulting in tradeoffs between investment in mate competition and somatic (anti-cancer) maintenance^28^. Sexually dimorphic or dichromatic species with extreme phenotypes, such as large and colorful ornaments or weapons, may have an increased risk of cancer^28^. This may be because selection for rapid cell growth in these tissues leads to the potential increased tumor growth as a byproduct. It may also be that there is selection for increased allocation of resources towards these costly sexual traits^77, 78^ at the expense of DNA repair and immune defenses^28^. However, even though testosterone in male red-legged partridges can increase the concentration of carotenoids, responsible for colorful traits, and testosterone suppresses the immune system, carotenoids also have immunoenhancing effects^79^. We found no significant difference in cancer prevalence in relation to sexual dimorphism and dichromatism. When factoring in both hue (the dominant wavelength of color) and brightness (the intensity of color), the degree of sexual dichromatism showed no significant correlation with neoplasia or malignancy prevalence. While most males tend to be larger than the females, that is not always the case, especially within birds of prey^80^. When examining the degree of sexual size dimorphism, we found no significant difference in cancer prevalence and differing sizes between sexes. This means that sexually dimorphic birds who spend time and energy in creating colorful plumage or larger body parts do not seem to pay a cost in terms of cancer susceptibility. It is possible that the birds in our study did not experience such tradeoffs because under human care they may have high energy budgets that allow them to invest both in sexually selected traits as well as in somatic maintenance in the form of cancer suppression. The same might not be the case for wild birds who are under greater energetic constraints and might therefore be more likely to experience tradeoffs.

### Do female birds have higher cancer prevalence than male birds?

Cancer rates in most other species, including humans, are biased toward males^16^. Current theory states that the double X chromosome found in females may offer some cancer protection^16^. For example, if the X chromosome carries a cancer-inducing mutation, the extra X chromosome present in females may carry a non-deleterious variant of the allele, whereas males (XY) without the extra X chromosome would not have this protective variant. In alignment with the two-X chromosome theory of cancer protection, previous work has shown that female birds (ZW) have more neoplasms than male birds (ZZ), but this was not validated statistically with sex-specific neoplasia prevalence^2^. We found that females do not have significantly different neoplasia prevalence or malignant prevalence than male birds. This analysis was done excluding reproductive cancers because living in managed environments with controlled reproduction could be affecting the animals’ susceptibility to cancers of the reproductive system.

### Future directions

We constructed a large and high-quality dataset including not only a significantly larger number of life history variables for birds than previous studies, but also detailed necropsy information for a large number of individuals per species, allowing greater error reduction, the inclusion of potential covariant traits, as well as the ability to distinguish benign and malignant tumors. Still, our study does not have information about the exact tissue where neoplasms were found in every individual, and future studies would benefit from knowledge of the relationships between distinct cancer types and life history in birds. There may also be evolutionary mismatches between animals in zoological institutions and in the wild. For example, peregrines^81^ and 84% of the mammalian species analyzed by Tidière et al.^82^ lived longer in zoos than in the wild. However, no significant difference was found in the maximum lifespan of 6 families of birds under human care versus wild birds (16 species of Anatidae, 3 species of Ciconidae, 10 species of Accipitridae, 6 species of Gruidae, 7 species of Corvidae, 3 species Pelecanidae)^83^. Future studies using a larger dataset with tracked life history and cancer records for every individual and tissue from birds in zoological institutions and in the wild would be helpful to better understand the role of life-history traits in cancer susceptibility.

Recent studies have focused on the evolutionary history of specific oncogenes in birds^84^. Specifically, the expansion of an oncoprotein, Golgi phosphoprotein 3, may contribute to birds’ relatively lower cancer susceptibility^84^ compared to mammals^2, 17, 18^. Although Golgi phosphoprotein 3 has many functions, such as modulating the dynamics of adhesion^85^ and regulating the function of mitochondria^86^, its exact molecular association with cancer suppression is not entirely clear^84^. Future work could examine the possible variation in the number of oncogenes and tumor-suppressors across bird species to identify how they are linked with cancer susceptibility and large clutch/litter size, and whether this correlation occurs in wild animals or is an artifact of domestication and artificial selection.

Several ecological factors may be driving many of the cancers in birds in our dataset. Previous work in chickens has shown that spontaneous and experimental infection with toxoplasma leads to the development of glioma-like tumors^87, 88^. Tumors were also detected in 25 out of 1669 free-living birds in the area of Chernobyl and were positively correlated with exposure to radiation^89^. To assess whether infections, radiation, or even nutritional factors, such as and carnivorous diets^90^, are associated with the malignancies and neoplasms of birds in our dataset, a systematic analysis of the carcinogens that these birds may be exposed to in managed settings would be necessary. This would also inform us about potential mechanisms that protect birds from radiation-induced DNA damage^91^, as well as associations between unpredictable environments and fast life history strategies (e.g., production of more offspring)^92^ that explain cancer susceptibility across species.

## Conclusions

We explored cancer prevalence across 108 managed species of birds. We found that among the examined life history factors, only clutch size was correlated (positively) with malignancy prevalence and neoplasia prevalence. Our findings are consistent with previous work which looked across 37 species of mammals in managed environments, finding that species with larger litter sizes were more vulnerable to cancer^35^. Further work is necessary, however, to examine whether these patterns hold up in wild and free-ranging populations.

## Supporting information

Life history and cancer dataset used in this study

## Acknowledgements

Thanks to all of the pathologists, veterinarians, and staff at the zoos, aquariums, and private veterinary centers for contributing to the data collection by diagnosing malignancy prevalence and neoplasia prevalence. Specifically, we would like to acknowledge the following institutions: Akron Zoo, Atlanta Zoo, Audubon Nature Institute, Bergen County Zoo, Buffalo Zoo, Capron Park Zoo, Central Florida Zoo, Dallas Zoo, El Paso Zoo, Elmwood Park Zoo, Fort Worth Zoo, Gladys Porter Zoo, Greensboro Science Center, Greenville Zoo, Henry Doorly Zoo, Utah’s Hogle Zoo, Jacksonville Zoo, John Ball Zoo, Los Angeles Zoo, Louisville Zoo, Miami Zoo, Northwest ZooPath, Oakland Zoo, Oklahoma City Zoo, Philadelphia Zoo, Phoenix Zoo, Point Defiance Zoo, Pueblo Zoo, San Antonio Zoo, Santa Ana Zoo, Santa Barbara Zoo, Sedgwick County Zoo, Seneca Park Zoo, The Brevard Zoo, The Detroit Zoo, The Oregon Zoo, and Toledo Zoo. Thanks to Diego Mallo and Walker Mellon for help with the statistical analyses. This work was supported in part by NIH grants U54 CA217376, U2C CA233254, P01 CA91955, and R01 CA140657 as well as CDMRP Breast Cancer Research Program Award BC132057 and the Arizona Biomedical Research Commission grant ADHS18-198847. The findings, opinions and recommendations expressed here are those of the authors and not necessarily those of the universities where the research was performed or the National Institutes of Health.

## Author contributions

This work was part of J.D.’s Barrett’s Honors College thesis at Arizona State University. A.A., A.M.B., Z.C., E.G.D., T.M.H., V.K.H., and S.M.R. helped in the collection of neoplasia prevalence, malignancy prevalence, and life-history trait data across species. Z.C. initially analyzed the sex bias cancer data and created the sex bias regression figures. S.E.K. contributed to a later version of the manuscript, reanalyzed data, updated figures, and rewrote sections of the manuscript for final submission to the journal. S.A. contributed in making the age density analyses. A.M.B., A.A., and C.C.M. provided helpful discussions, comments, and guidance throughout the project. All authors commented on the final versions of the manuscript.

## Supplementary material

**Supplementary Figure 1.** The log_10_ of adult mass times lifespan is not correlated with neoplasia prevalence (A) or malignancy prevalence (B) across 57 bird species. The black lines show the phylogenetically-controlled linear regression of the log_10_ of adult mass times lifespan versus malignancy prevalence or neoplasia prevalence. Adult mass is measured in grams, whereas lifespan is measured in months. Different colors show the different order each species belongs to.

**Supplementary Figure 2.** No significant sex bias in neoplasia (A) or malignancy prevalence (A) **across 31 bird species.** Each dot in plot A shows the male neoplasia prevalence and female neoplasia prevalence of a species. Whereas each dot in plot B shows the male malignancy prevalence and female malignancy prevalence of a species.

**Supplementary Figure 3. Cancer deaths are not skewed towards old age.** Normalized frequency of a species’ age at death as a percentage of the species lifespan. Each density plot shows the necropsied individuals that had tumors (blue) and the necropsied individuals that did not have tumors (red). There are 1287 individuals in this distribution from which we have lifespan data.

**Supplementary Figure 4.** Larger clutch size is correlated with malignancy prevalence (B) but not neoplasia prevalence (A) across 45 bird species after removing domesticated and semi-domesticated species from the analyses. After controlling for species body mass, the positive correlation between clutch size and malignancy prevalence remains significant (*P*-value = 0.004; Table 2). Dot size shows the number of necropsies per species. Colors show the taxonomic order of each species. Black lines indicate the phylogenetically-controlled linear regression of the normalized values of clutch size versus malignancy prevalence or neoplasia prevalence.

**Supplementary Figure 5.** Pearson’s correlation matrix with four life history variables shared by 19 species in our dataset (log_10_ of adult mass, lifespan^0.425^, incubation length, –1 **clutch size^−0.1^**^25^).

## Supplementary data

Life history and cancer dataset used in this study.

**Supplementary Table 1.** Common species analyzed in Møller et al.^19^ and our study. Here we present species’ tumor incidence and number of records in Møller et al.^19^ versus neoplasia prevalence and number of necropsies in our study. We also present the *P-*values of Fisher’s exact test.

| **Species (common name)** | **Tumor incidence in Møller et al.'s study (# records)** | **Neoplasia prevalence in this study (# necropsies)** | ***P*-value** |
| --- | --- | --- | --- |
| *Columba livia* (rock pigeon) | 0% (3) | 1.8% (55) | 1 |
| *Anas acuta* (northern pintail) | 0% (3) | 4.3% (23) | 1 |
| *Milvus milvus* (red kite) | 0% (3) | 0% (36) | 1 |
| *Crex crex* (corncrake) | 0% (4) | 0% (47) | 1 |
| *Fringilla coelebs* (chaffinch) | 0% (213) | 0% (45) | 1 |
| *Anas platyrhynchos* (mallard duck) | 4.7% (21) | 21.2% (33) | 0.13 |

