## Supplementary figures and images for "Life history and cancer in birds: clutch size predicts cancer"

### Supplementary Figure 1

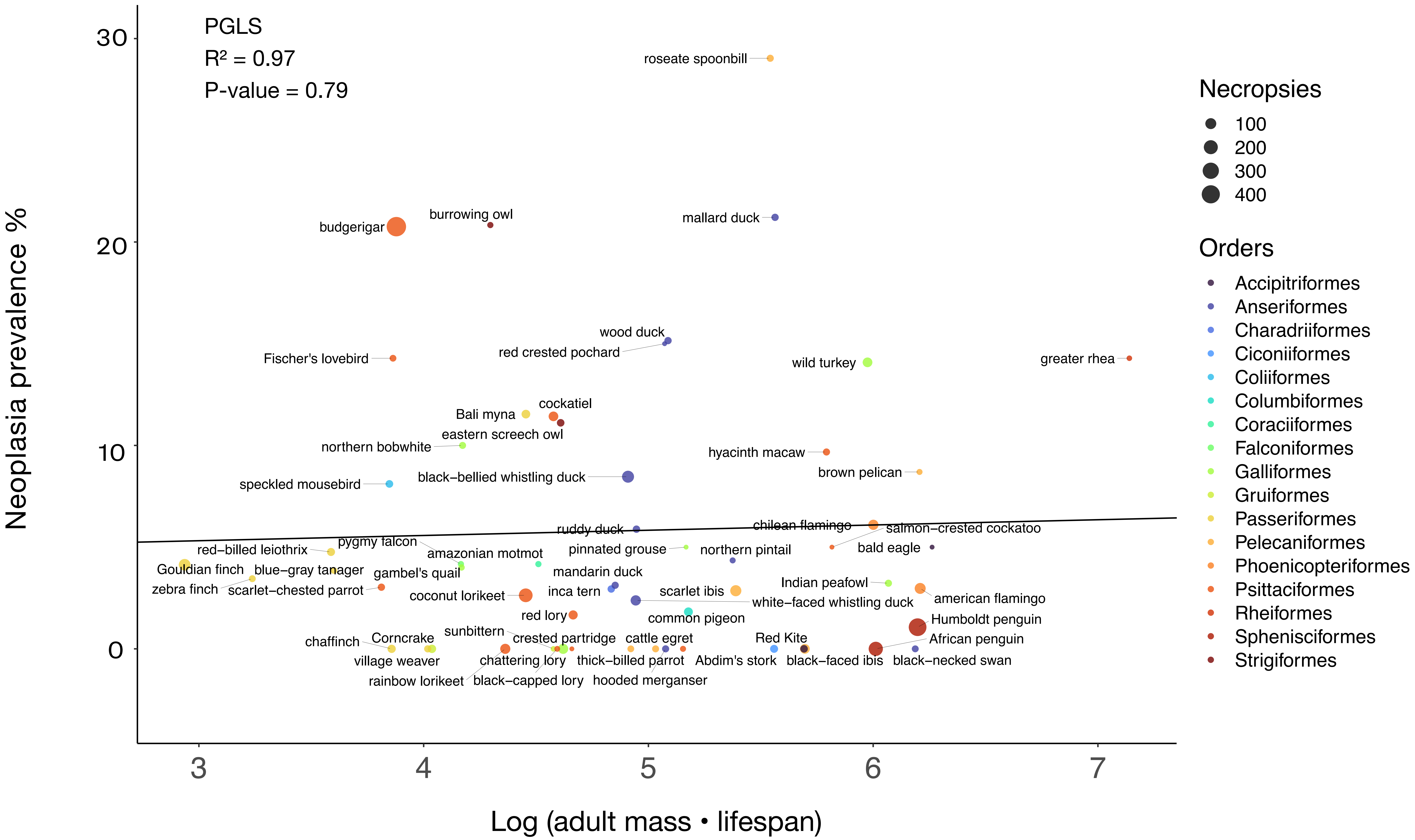

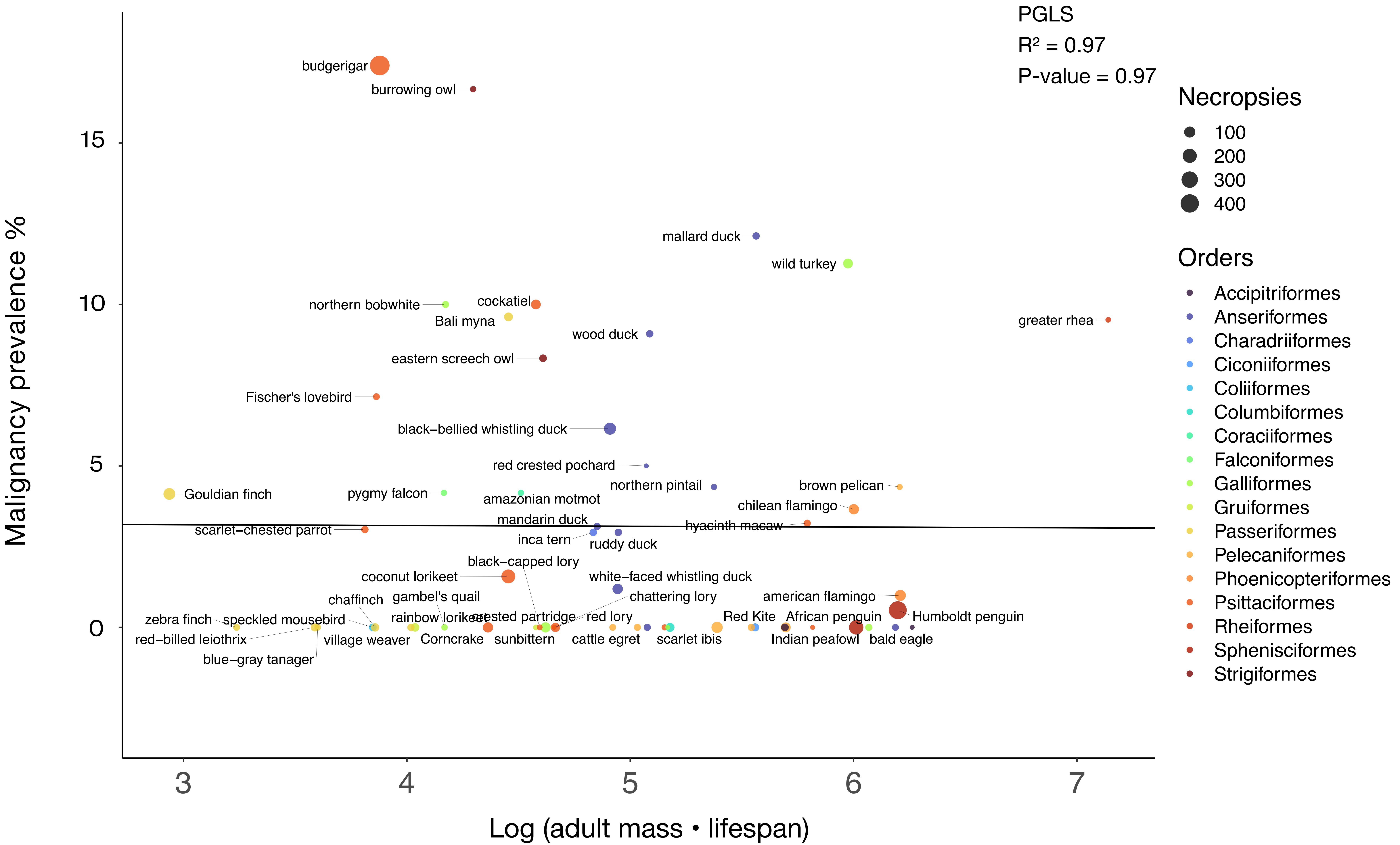

### Supplementary Figure 2

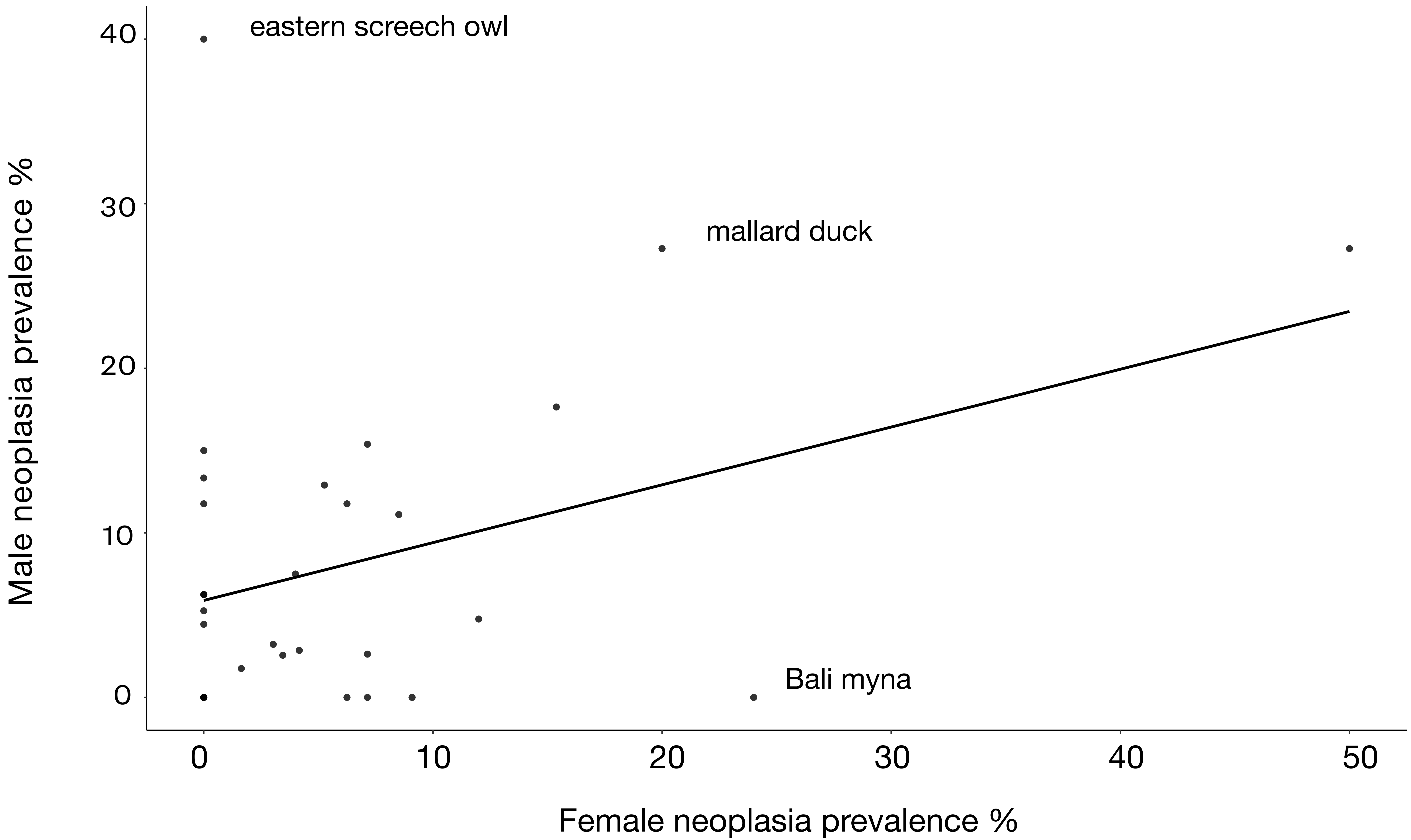

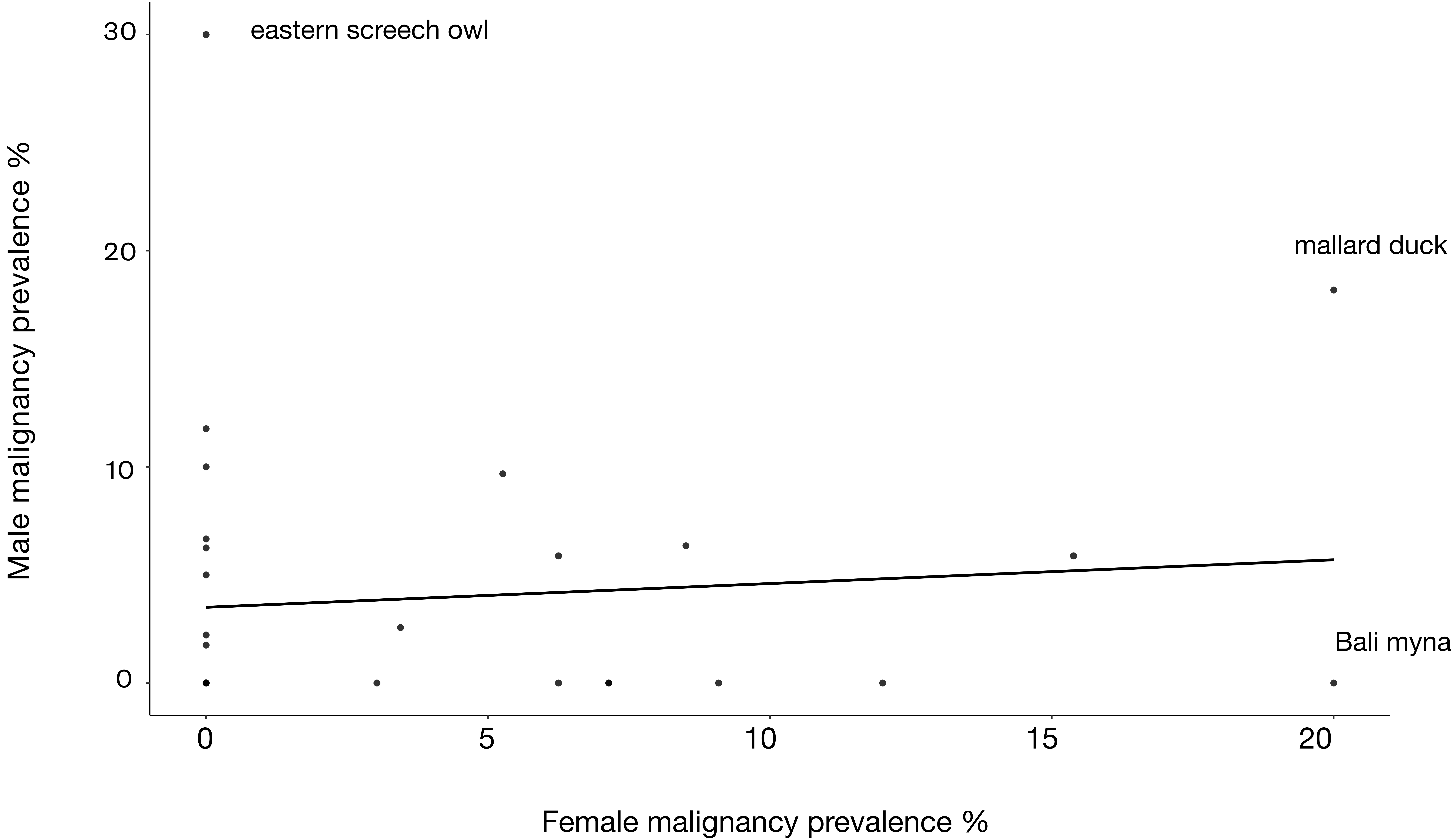

### Supplementary Figure 3

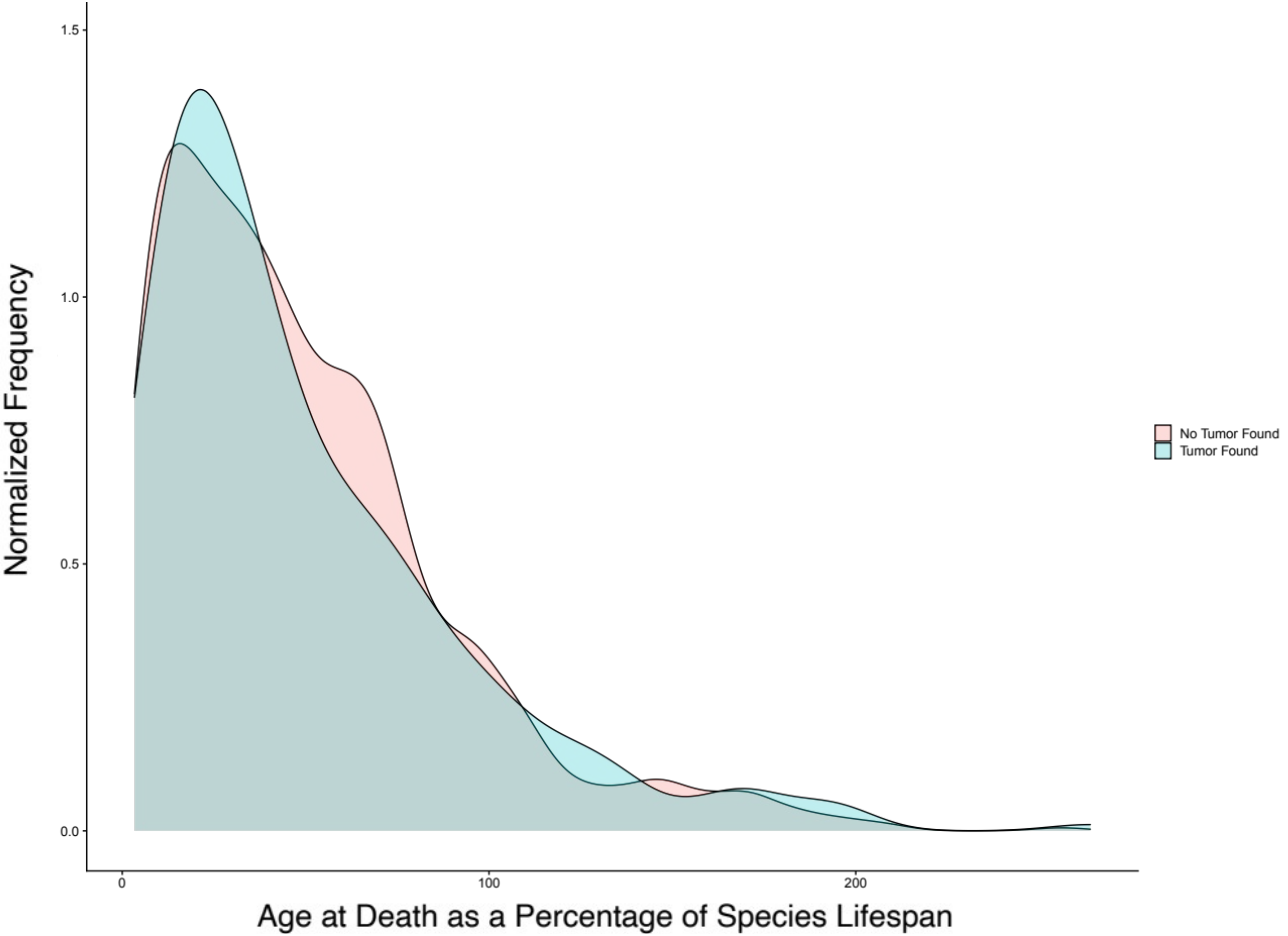

### Supplementary Figure 4

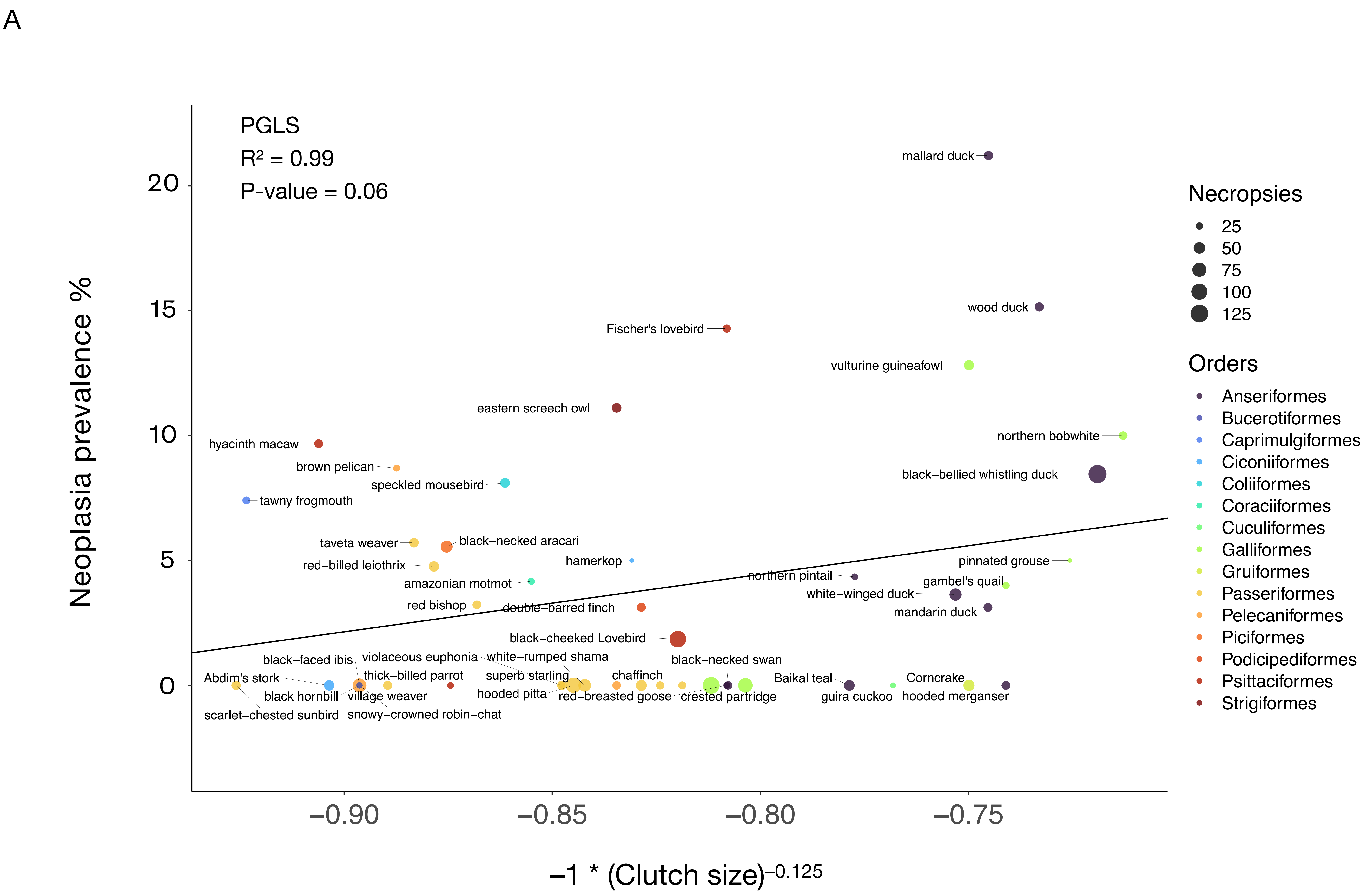

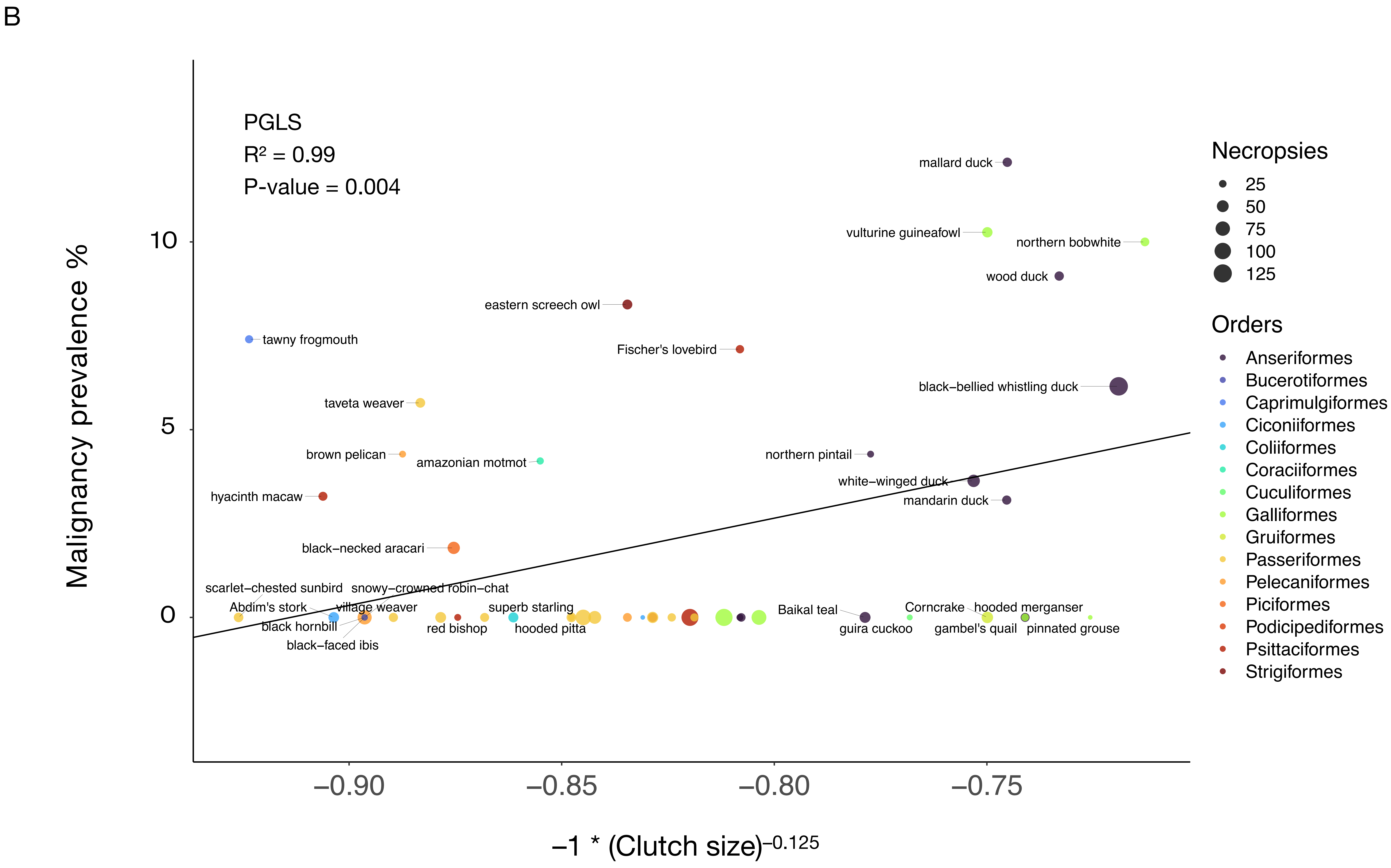

### Supplementary Figure 5

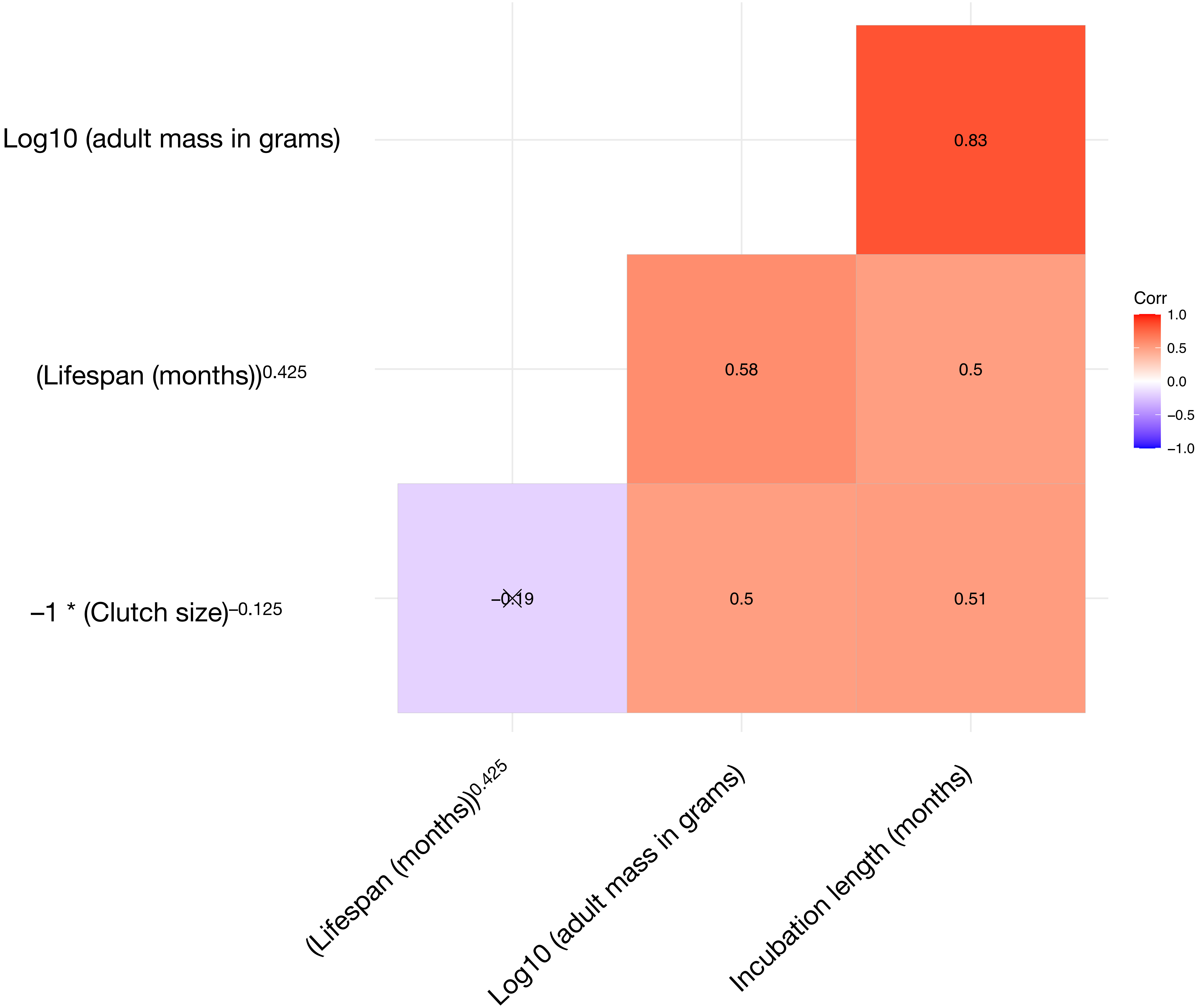
